# Tight junctions regulate lumen morphology via hydrostatic pressure and junctional tension

**DOI:** 10.1101/2023.05.23.541893

**Authors:** Markus Mukenhirn, Chen-Ho Wang, Tristan Guyomar, Matthew J. Bovyn, Michael F. Staddon, Riccardo Maraspini, Linjie Lu, Cecilie Martin-Lemaitre, Masaki Sano, Tetsuya Hiraiwa, Daniel Riveline, Alf Honigmann

## Abstract

Formation of fluid filled lumen by epithelial tissues is a fundamental process for organ development. How epithelial cells regulate the hydraulic and cortical forces to control lumen morphology is not completely understood. Here, we quantified the mechanical role of tight junctions in lumen formation using genetically modified MDCKII cysts. We found that the paracellular ion barrier formed by claudin receptors is not required for hydraulic inflation of lumen. However, depletion of the zonula occludens scaffold resulted in lumen collapse and folding of apical membranes. Combining quantitative measurements and perturbations of hydrostatic lumen pressure and junctional tension with modelling, we were able to predict lumen morphologies from the pressure-tension force balance. We found that in MDCK tissue the tight junction promotes formation of spherical lumen by decreasing cortical tension via inhibition of myosin. In addition, we found that the apical surface area of cells is largely uncoupled from lumen volume changes, suggesting that excess apical area contributes to lumen opening in the low-pressure regime. Overall, our findings provide a mechanical understanding of how epithelial cells use tight junctions to modulate tissue and lumen shape.

## Introduction

Luminal cavities are key structural components of organs, functioning as compartments for fluid transport, nutrient absorption or waste secretion. Establishment of lumen during development occurs via different morphogenetic mechanisms^1^. Lumen that form between existing cell contacts such as acini, blastocytes or canaliculi require mechanisms that enable expansion of a fluid filled cavity against the resistance of the surrounding cells.

On the molecular level lumen formation is often coupled to the establishment of apical-basal polarity. In particular, formation of an apical membrane with a distinct composition that excludes cell-cell adhesion receptors and enriches ion transporters is important for lumen formation^2^. Molecular activities such as ion pumping, and acto-myosin constriction have been related to the mechanical forces that are required to open a luminal cavity between cells^3, 4^. Current physical models explain lumen shapes by considering osmotic, hydraulic and cortical forces with a special emphasis on luminal pressure^3, 4^. Lumen inflation results from water influx that arises from ion gradients and hydrostatic pressure. However luminal shape is a result of the competition between hydrostatic pressure and mechanical properties of the surrounding cells. In addition to this hydraulic based mechanism, it has been suggested that lumen can also grow in a pressure independent way via forces that arise from formation and maintenance of a constant apical membrane area per cell^5^. Despite this progress, it remains challenging to link the control of mechanical forces that govern lumen growth and shape to the molecular regulation that happens on the cellular and tissue level.

Here, we investigated the role of epithelial tight junctions in lumen growth and shape using MDCKII cysts, an organotypic cell-culture model for lumenogenesis. Tight junctions control the paracellular flow of ions and water across the tissue by forming a trans-epithelial diffusion barrier via claudin strands^6^. Knock-out of claudins in MDCKII cells (CLDN-KO) results in complete loss of TJ-strands and CLDN-KO tissue is permeable for small molecules such as ions but remains impermeable to large macro-molecules^7, 8^. Interestingly, the cytoplasmic TJ-scaffold of ZO proteins remains largely unaffected by claudin depletion. The ZO scaffold of tight junctions has been shown to regulate the structure and activity of the actomyosin cortex^9–11^. Depletion of ZO1/ZO2 in MDCKII cells causes complete loss of all tight junction proteins from the sub-apical belt including claudins^7, 12, 13^. In addition, the actin cytoskeleton is drastically remodeled^9–11^. Accordingly, ZO-KO tissue is highly permeable for small and large molecules^7, 14^. Since the control of both transepithelial permeability and cytoskeletal tension are key mechanical parameters for lumen formation, tight junctions are expected to be an important molecular regulator of lumen morphology^7, 9–11, 14^. Here, we studied lumen formation of genome edited MDCKII cysts mutated for key tight junction proteins (WT, CLDN-KO, ZO-KO) using quantitative microscopy and mechanical measurements of lumen pressure and junctional tension.

Surprisingly, we found that the opening of the transepithelial barrier to ions, using MDCKII cells depleted of all expressed claudins (CLDN-KO,^7, 8^), does not result in a decrease of hydrostatic lumen pressure. Despite high ion leakage, lumen pressure is maintained close to wild-type levels and lumen expand to a round shape similar to wild-type tissue. This suggests that cells can compensate para-cellular leakage by increased ion pumping. In contrast, depletion of the cytoplasmic tight scaffold (ZO-KO)^13^ resulted in lumen volume collapse accompanied by a loss of hydrostatic pressure and a strong increase in junctional tension. Interestingly, the surface area of the collapsed lumen remained similar to WT-tissue causing strong deformations of the apical membrane, suggesting that the apical membrane area is largely independent of hydrostatic lumen pressure and cortical tension. Using a mechanical force balance model with a constrained apical area, we predicted lumen morphologies as a function of lumen pressure and junction cortical tension and found strong agreements with experimental phenotypes. Finally, we used perturbation of lumen pressure and junctional tension to verify the mechanical model using specific inhibitors and a pipette-mediated inflation assay. Taken together, our results revealed that an important function of tight junctions is to reduce the mechanical tension of cells, which enables lumen expansion at low hydrostatic pressure. We envision that the regulation of the pressure-tension force balance by tight junctions is an important control node that allows tuning tissue and lumen shapes during morphogenesis.

## Results

### Tight junction (TJ) mutants used and their properties

To investigate the function of TJs in lumen formation (Fig. 1A), we used genome edited MDCKII cells lacking either the set of both cytoplasmic tight junction scaffold proteins ZO1 and ZO2 (ZO-KO,^13^) or the set of five major claudins expressed in MDCKs (CLDN-KO,^7^ Fig. 1B). ZO1 and ZO2 as well as claudins are essential for the assembly of tight junction strands and formation of a tissue permeability barrier. In addition, ZO proteins are involved in regulation of cell mechanics via interactions with the acto-myosin cortex^6, 7, 13, 15^.

**Figure 1:**
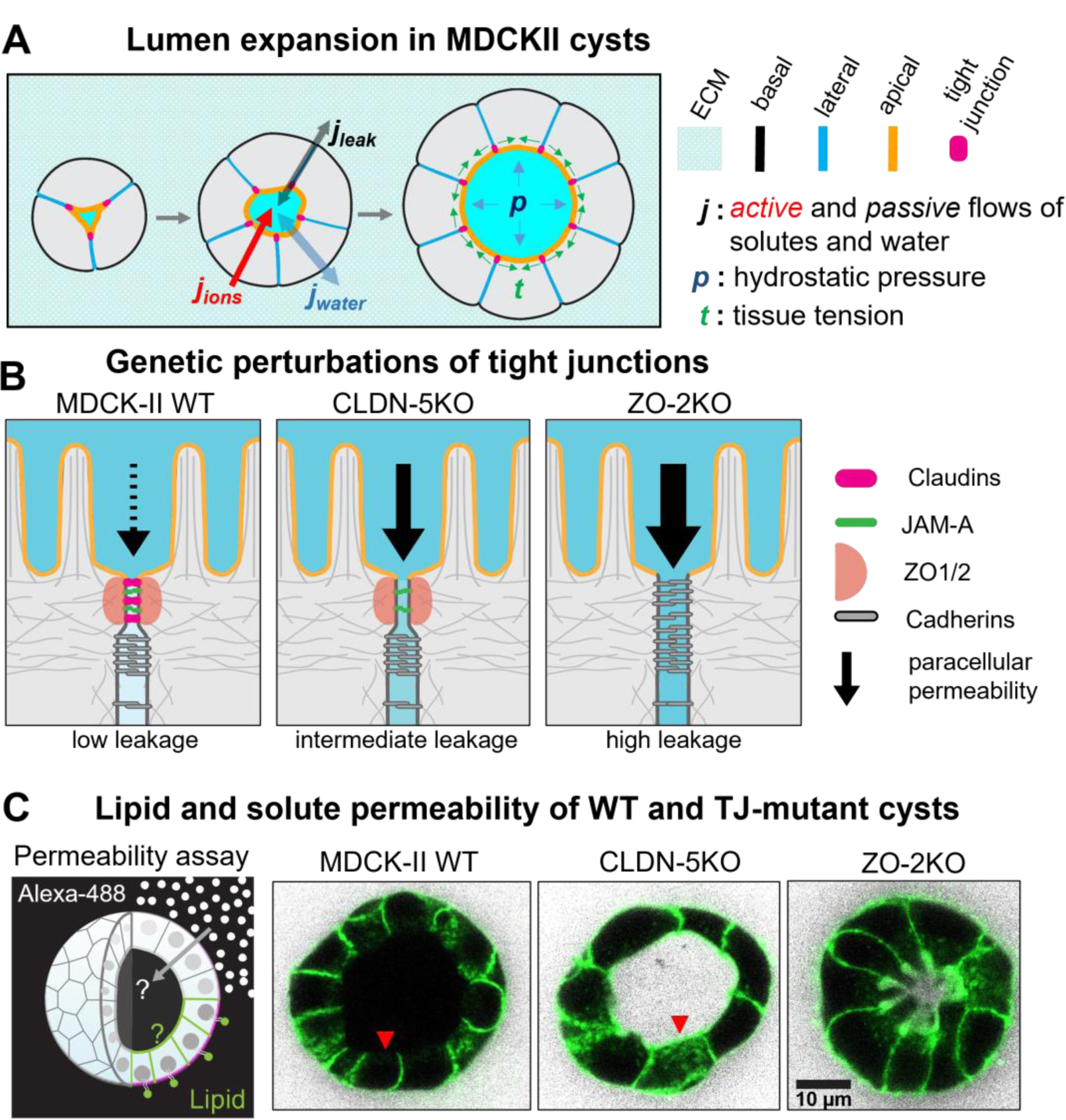
Lumen formation in MDCKII cysts and tight junction mutants used in this study. (A) Scheme of lumen expansion in MDCKII cysts. Polarized membrane domains and tight junctions are highlighted (right panel). Cysts are growing over time due to cell divisions. Osmotic gradients due to the balance of active ion transport into the lumen and passive para-cellular leaks result in water influx and establishment of hydrostatic lumen pressure. Hydrostatic lumen pressure is balanced by cortex tension of the apical-junctional surface. (B) Tight junction mutants used in this study. MDCKII-WT tissue produces tight junctions that have very low permeability for ions and water. CLDN-5KO tissue is permeable for small molecules and ZO-2KO tissue is permeable for small-and macromolecules. (C) Permeability assay in MDCKII cysts for small molecules (Alexa-488) and a lipid tracer (Cell-Mask). Shown are midplane cross-section of WT and TJ-mutant cysts, white is the small molecule tracer and green the lipid tracer. WT cysts are tight for both molecules. CLDN-KO and ZO-KO cysts are permeable for both molecules. However, CLDN-KO cysts produce a large lumen while ZO-KO cysts produce small, convoluted lumen. Arrows indicate apical membranes, which are not accessible for the lipid tracer in WT cysts but in CLDN-5KO and ZO-2KO the lipid tracer can cross the junctional barrier.

Previous studies of these TJ-mutant cell lines have been performed using mostly 2D tissue culture. To study epithelial barrier formation in 3D, we used MDCKII cysts embedded in Matrigel and determined the permeability of a small molecular tracer (Alexa488) and a lipid dye to the luminal side of the tissue. In WT cysts, the lipid dye and the soluble small molecule tracer were excluded from the apical membrane (Fig. 1C). In contrast, lumen of cysts grown from CLDN-KO and ZO-KO cells were permeable to the tracer as well as the lipid dye (Fig. 1C). We noted that CLDN-KO cysts were able to form a large, inflated lumen despite being permissive to small solutes, whereas ZO-KO cysts had significantly smaller and more convoluted lumen shapes (Fig. 1C). In line with previous reports^7, 13^, these results demonstrate that loss of tight junctions causes increased epithelial barrier permeability and that loss of ZO1/ZO2 altered lumen morphology.

### Quantification of cyst and lumen morphology of TJ mutants

To quantify the impact of TJs on lumen formation, we compared changes in lumen and tissue morphology of WT, ZO-KO and CLDN-KO during cystogenesis in Matrigel (Fig. 2A). Immunofluorescence staining (IF) was performed on cysts at different stages of cyst development and a 3D image segmentation and analysis pipeline was established to acquire morphometric parameters of lumen and cysts.

**Figure 2:**
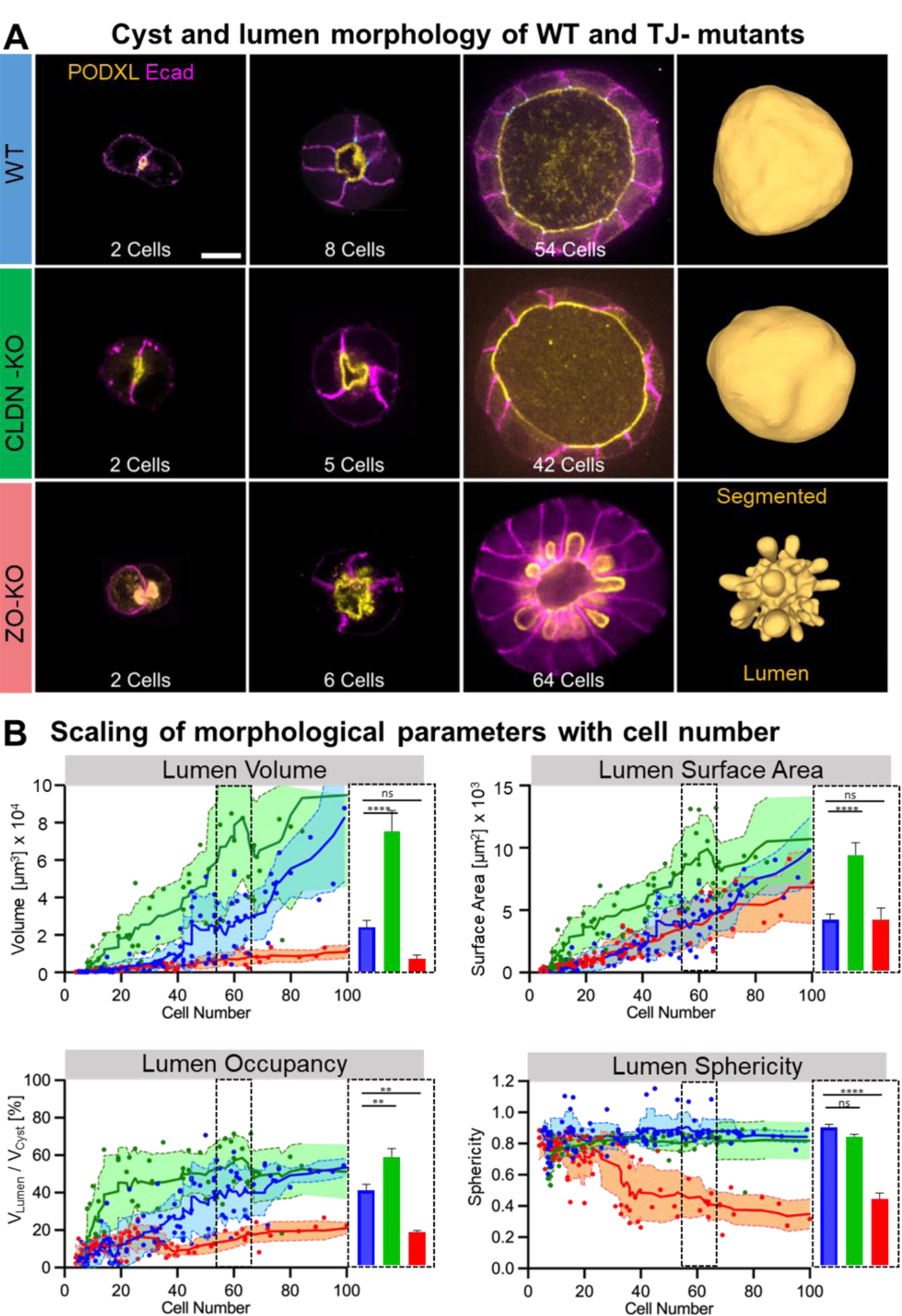
Lumen morphological differences between WT and TJ-Mutants cell lines. (A) Midplane cross-section of WT and TJ-mutant cysts as a function of cell number (indicated at the bottom of the panel). Lateral membranes are stained by E-cadherin (magenta) and apical membranes by Podocalyxin (yellow). The right panels show 3D segmentation of the lumen at the later stage. WT and CLDN-KO cyst form large spherical lumen, ZO-KO cysts form smaller, convoluted lumen with apical membranes curving into the cell bodies. Scalebar is 10µm. (B) Quantification of lumen volume, surface area, sphericity and occupancy based on 3D segmentation of lumen and cyst surfaces of WT (n =114), CLDN-KO (n=64) and ZO-KO cysts (n=88) as a function of the total cell number per cyst. Bar graphs indicate quantification of lumen volume, surface area, sphericity and occupancy for 55 – 65 cells (dashed box) and WT (n =15), CLDN-KO (n=6) and ZO-KO cysts (n=9), Oneway-Anova was used for statistical analysis.

Overall, we observed that CLDN-KO and ZO-KO cells formed cysts with a single lumen and polarity markers such as podocalyxin (apical) or E-Cadherin (lateral) that were correctly polarized as in WT tissue (Fig. 2A). Furthermore, lumen morphology at initial cyst stages (2-6 cells) was comparable between WT, CLDN-KO and ZO-KO cysts (Fig. 2A, B). Interestingly, at later stages of cyst development (>20 cells), clear differences in lumen morphology emerged: While both WT and CLDN-KO cysts developed large spherical lumen, ZO-KO cysts failed to inflate their lumen, which resulted in intricately folded apical membranes often bending into the cell bodies (see Video S1).

Quantification of lumen volume and apical surface area as a function of cell number revealed that on average lumen volume of WT and CLDN-KO cysts increased monotonically with cell number to a value of 8 × 10^4^ µm^3^ at 100 cells, with CLDN-KO lumen showing slightly faster growth than WT (Fig. 2B). In contrast, the lumen volume of ZO-KO cysts inflated significantly slower and remained below 1 × 10^4^ µm^3^ even at 100 cells. Interestingly, despite ZO-KO cysts having a strongly reduced lumen volume, the apical surface area of lumen increased linearly with cell number with the same slope as WT cysts. Thus, the apical surface area per cell was constant over time and comparable between WT and ZO-KO cells. This indicates that lumen volume and surface are not coupled, e.g., cells do not adapt their apical area according to changes in lumen volume.

To quantify the deviation from a spherical shape we calculated the sphericity of the lumen over time. After an initially irregular shape, sphericity of WT and CLDN-KO quickly reached values close to a perfect sphere around the 10 to 20 cell stage. In contrast, sphericity of ZO-KO lumen dropped as a function of cell number, showing that the lumen surface to volume ratio increased as more and more cells compete for the relatively small lumen volume over time. To determine how the lumen volume grows in relation to the whole cyst volume we calculated the lumen occupancy, i.e., the ratio of lumen to cyst volume. Lumen occupancy saturated around 50% for WT and CLDN-KO cysts. However, the latter reached saturation already at the 20-cell stage while WT cyst took 60-80 cells to reach a similar lumen occupancy, which indicates that lumen expansion in CLDN-KO cysts happens faster than WT despite higher trans-epithelial leakage. ZO-KO lumen occupancy was significantly lower than WT and saturated around 20%. The low lumen occupancy of ZO-KO cysts went along with a significant thickening of the epithelial cell layer with cells having columnar shapes compared to the cuboidal shapes of WT cysts (see Fig. 2A).

Taken together, the morphometric analysis revealed that perturbing TJs via claudin or ZO depletions has differential effects on lumen morphogenesis. First, lumen inflation is strongly reduced by ZO-KO, while CLDN-KO cysts inflate spherical lumen like WT cysts. Second, the apical area of cells is largely constant and uncoupled from the collapse of lumen volume in ZO-KO tissue. Since both mutations produce a strong increase in trans-epithelial ion permeability compared to WT tissue, it is interesting that CLDN-KO cysts are able to inflate large spherical lumen whereas ZO-KO depletion caused a deflated lumen. To address this difference, we turned to quantification of mechanical parameters starting by designing a new method to measure hydrostatic pressure of lumen.

### Quantification of hydrostatic lumen pressure by lumen drainage

In many epithelial tissues lumen inflation has been linked to establishment of hydraulic forces^16^. Apical secretion of chloride ions has been suggested to drive osmotic water influx into the lumen of MDCK cysts, producing a hydrostatic pressure difference that inflates the lumen^17^. To test whether the morphological difference in lumen morphology between WT, CLDN-KO and ZO-KO cysts are related to changes in hydrostatic pressure, we designed a lumen drainage experiment.

To measure the hydrostatic pressure difference between the lumen and the extracellular fluid, we used a two-photon laser to cut the cyst open via a short line scan through a single cell (Fig. 3A). Cyst opening resulted in subsequent collapse of the lumen volume, which we quantified by segmenting the whole cysts and lumen from 3D confocal stacks over time (Fig. 3A, Video S2). We found that the lumen volume of WT and CLDN-KO cysts decreased with an exponential-like kinetic to ≈ 20 % after 300 s, while ZO-KO showed significantly less decrease to ≈ 70 % of its initial volume (Fig. 3D).

**Figure 3:**
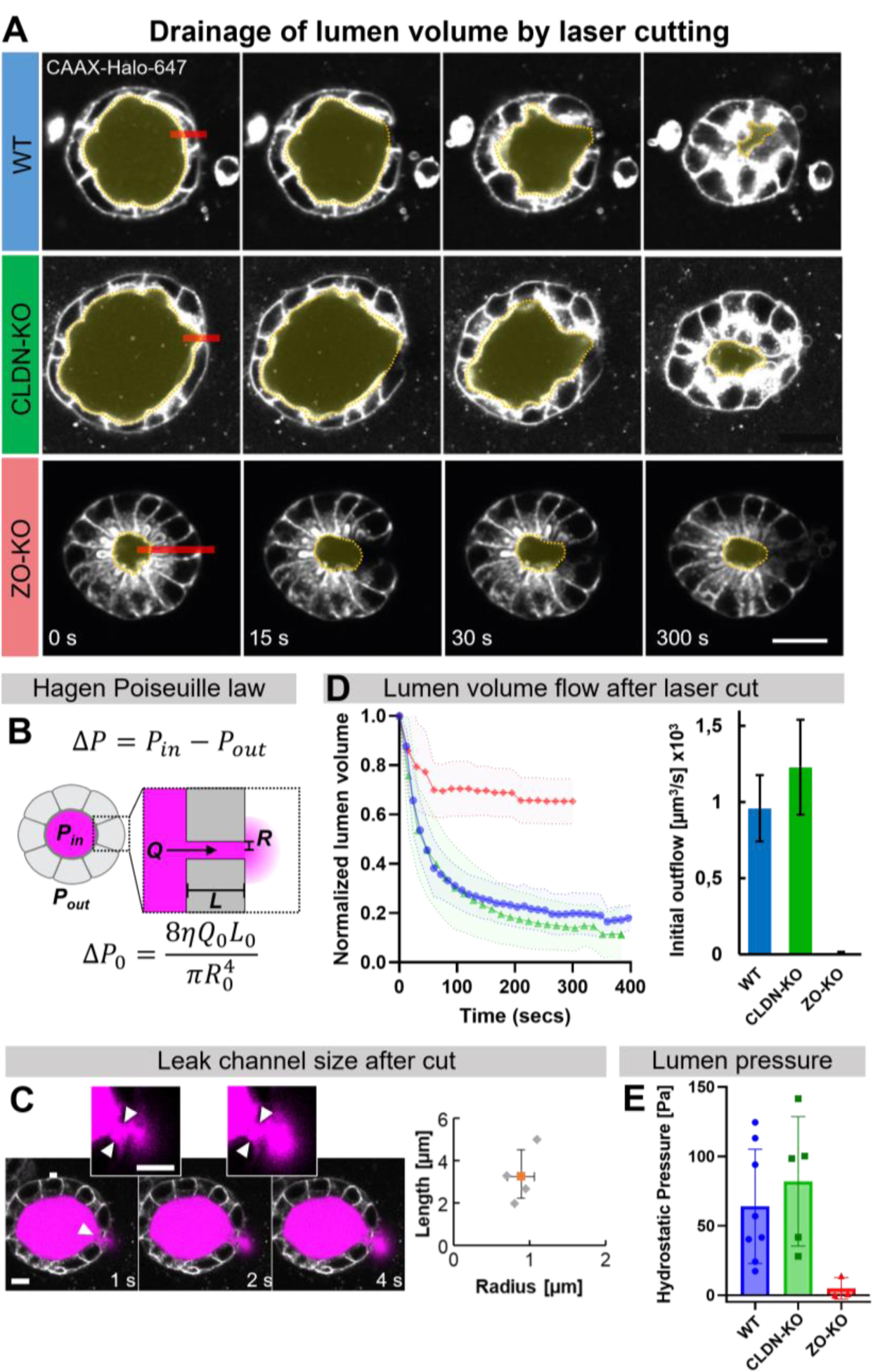
Influence of TJ-mutations on hydrostatic lumen pressure in MDCKII cysts. (A) Lumen volume drainage MDCKII TJ mutant cysts by laser cutting. Midplane cross-section of cysts before and after laser cutting. Position of cuts are indicated by red lines. The lumen of WT and CLDN-KO cysts quickly collapsed after the cut, while the lumen of ZO-KO cyst remained relatively constant. Scalebar is 20µm. See supplemental movie-1 for an example of lumen drainage. (B) Hagen-Poiseuille law used to determine initial hydrostatic pressure in the lumen: *ΔP* hydrostatic pressure difference, *Q* fluid flow, *L* and *R* channel length and radius, *η* fluid viscosity. Lumen fluid viscosity was measured using FCS (Suppl. Fig. 1). (C) Geometry of the leak channel after laser cutting was determined by imaging the exit of the lumen fluid, which was stained by expression of Dendra2-CD164 (magenta). Scalebar 10 µm. (D) Lumen volume change after laser cutting determined from 3D segmentation of lumen over time. Values are normalized to the initial lumen volume before the cut. Right plot shows mean initial flow rate with standard error, calculated from the initial slope. WT (n=12), CLDN-KO (n=6) and ZO-KO cysts (n=5). (E) Hydrostatic lumen pressure of MDKC-II mutants. WT and CLND-KO have a lumen pressure of ≈ 65 Pa and ≈ 85 Pa while ZO-KO cysts are not able to build up significant pressure.

We quantified the hydrostatic lumen pressure *ΔP* using the Hagen-Poiseuille law, which describes a flow *Q* of a fluid with viscosity *η* through a channel with radius *R* and length *L* due to a hydrostatic pressure difference *ΔP* (Fig 3B and Material and Methods). To measure the fluid flow together with the exit channel size, we expressed a fluorescently tagged, apically secreted sialomucin (Dendra2-CD164) in WT cysts, which provided a strong staining of the luminal fluid. This allowed us to visualize the fluid outflow from the lumen after laser-cutting the cyst and also to determine the average length and radius of the exit channel assumed to be cylindrical (Fig. 3C, Video S3, and Supplementary Materials). The outflow was quantified via segmentation of luminal volume change from 3D confocal stacks over time (Fig. 3A). The flow rate *Q* of luminal fluid was obtained from the initial change of lumen volume after cyst cutting (Fig. 3D).

Finally, to determine the viscosity of the luminal fluid, we measured the diffusion coefficient of Dendra2-CD164 in the lumen and in PBS buffer using fluorescence correlation spectroscopy. Lumen fluid viscosity was ≈ 5 times higher than water (Suppl. Fig. 1). Based on these four measured quantities (flow, radius, length and viscosity), we calculated the average hydrostatic lumen pressure to be ΔP_WT_ = 65 ± 15 Pa, ΔP_CLDN-KO_ = 84 ± 21 Pa and ΔP_ZO-KO_ = 0.5 ± 0.5 Pa (Fig. 3E).

Taken together, the hydrostatic pressure measurements revealed that WT cysts build up hydrostatic pressure of ≈ 65 Pa to inflate a spherical lumen, which is close to previously reported values^18^. In addition, despite CLDN-KO cyst being permeable for small molecules, they build up even higher lumen hydrostatic pressure than WT cysts ≈ 80 Pa. In contrast, ZO-KO cyst do not build up a significant hydrostatic pressure difference, which may explain why their lumen are collapsed. To understand how the changes in hydrostatic pressure are related to the morphological changes of the lumen, we next determined the corresponding changes of junctional tension of the TJ mutants.

### Tight junction mutations increase apical-junctional tension via myosin activation

To determine the influence of TJ-mutations on mechanical tissue tension, we quantified endogenous myosin-IIa distributions and performed laser ablation experiments. To visualize myosin distribution, we tagged endogenous Myosin-IIa with mNeonGreen in WT, CLDN-KO and ZO-KO cells using CRISPR/Cas genome editing. We observed that in WT monolayers Myo-IIa was mostly cytoplasmic, while in CLDN-KO and ZO-KO Myo-IIa was clearly enriched at the apical junctional complex (Fig. 4A). Quantification of the enrichment of Myo-IIa at the junction with respect to the cytoplasm, i.e., the junction to cytoplasm ratio (JCR), showed that myosin in WT tissue was only slightly enriched at the junction JCR ≈ 1, while in CLDN-KO monolayers myosin was strongly enriched at the TJ with JCR ≈ 5. ZO-KO showed an even stronger enrichment with JCR ≈ 7 (Fig. 4B).

**Figure 4:**
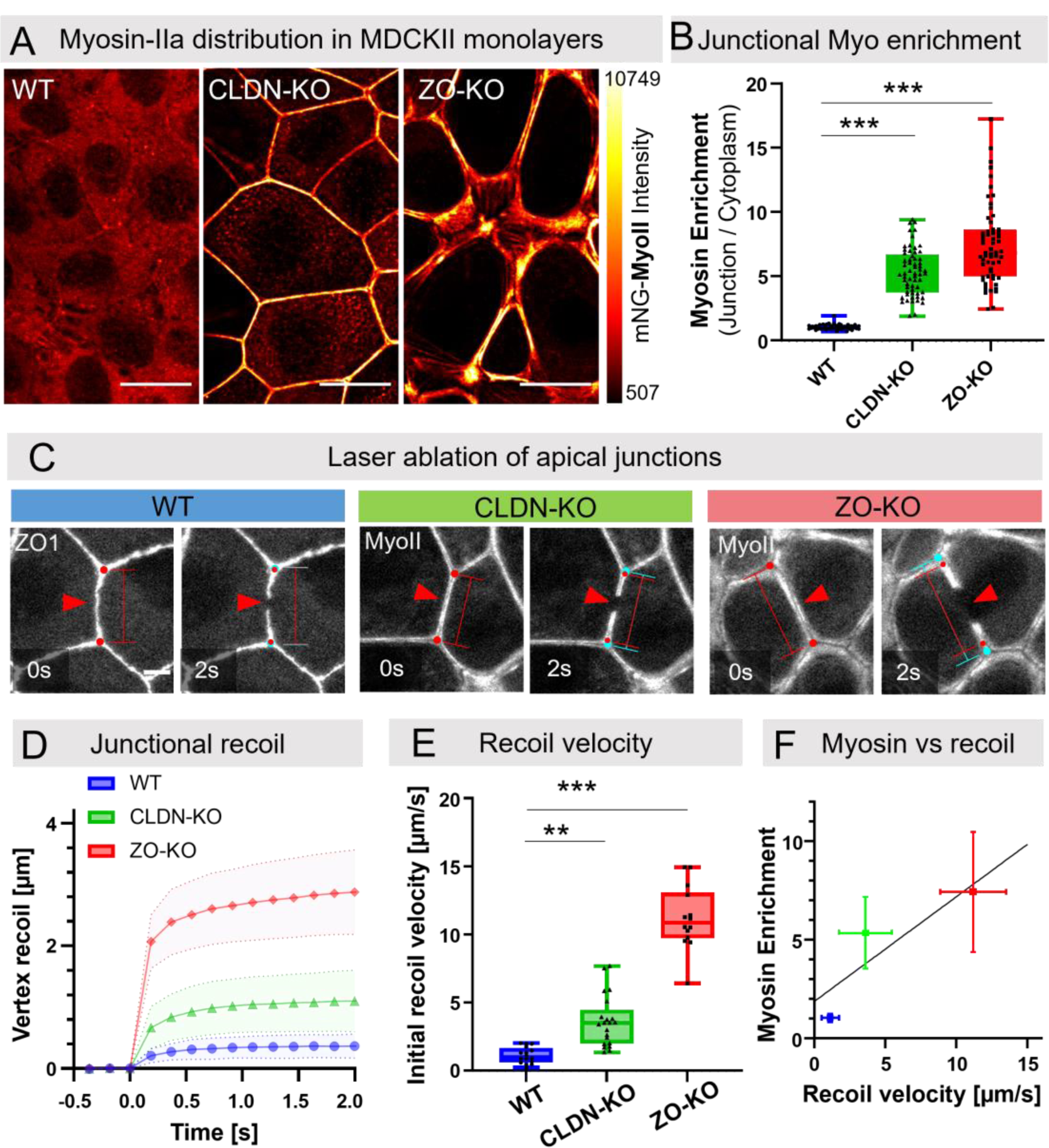
Quantification of junctional tension of MDCKII mutants. (A) Distribution of endogenous myosin-IIa tagged with mNeon in MDCK-II monolayers. In WT tissue myosin is predominantly in the cytoplasm with only minor punctae at cell junctions. CLDN-KO tissue showed significantly increased levels of myosin at the junctions and ZO-KO monolayer had strong accumulation of thick myosin bundles along cell-cell junctions. Scalebar is 5µm. (B) Quantification of junctional myosin enrichment at cell-cell junctions versus cytoplasm. WT (n =50), CLDN-KO (n=61) and ZO-KO cysts (n=60). (C) Laser ablation of cell-cell junctions of MDCK-II mutants. Position of cut is indicated by red arrow. Junctional recoil distance was measured by tracking the position of tricellular vertices over time. (D) Dynamics of junctional recoil after laser ablation of MDCK-II WT and mutants. WT (n =16), CLDN-KO (n=21) and ZO-KO cysts (n=14). Laser cut was done at 0s. Initial distance before cut was subtracted to highlight the change due to recoil. (E) Mean initial recoil velocities of MDCK-II mutants show that WT tissue is under low mechanical tension. CLDN-KO tissue has increased tension and ZO-KO tissue builds up strong junction tension. Initial recoil velocities were determined from the slope of first data points after laser cutting. WT (n =16), CLDN-KO (n=21) and ZO-KO cysts (n=14). (F) Correlation between myosin junctional enrichment and initial recoil velocity with a linear fit of WT, CLDN-KO and ZO-KO data from B and E (R^2^= 0.46).

Increased junctional myosin levels suggest that cortical tension of the tight junction mutants is increased. In order to estimate junctional tension, we laser dissected apical junctions and analyzed initial recoil velocities (Fig. 4C, D, E). WT tissue showed little recoil with a velocity of v = 1.1 μm/s, indicating that the junctions are not under significant tension and the tissue is in a relaxed state. In contrast, we found that CLDN-KO recoil was 3x faster (v = 3.6 μm/s) and ZO-KO even 10x faster (v =11 μm/s) compared to WT. Relating the junctional myosin levels to the junctional recoil velocities showed a strong positive correlation (Fig. 4F), which suggests that depletion of CLDN and even more so of ZO proteins increases junctional tension via local myosin accumulation.

Taken together, myosin localization and junctional recoil quantification confirmed that tight junctions play an important role in regulating junctional tension in MDCK tissue. ZO proteins and to a lesser extent claudins affect junctional myosin levels and as a consequence lower mechanical tension at the apical junctional complex, effectively making the tissue softer. These results are in line with previous observations that loss of ZO1/ZO2 and loss of claudins are linked to myosin accumulation at the apical junction and increased tissue tension^9–11^. Increased junctional tension has also been linked to apical constriction of cells, which can induce folding of tissues^19^. In the context of lumen formation, apical constriction of cells is expected to counteract the hydrostatic pressure of the lumen (Fig. 1A). Therefore, the combined loss of hydrostatic pressure together with the increased apical-junctional tension in ZO-KO cyst may explain the collapsed lumen volume and the folding of apical membranes.

The combined pressure and tension results suggest that tight junctions promote lumen inflation by two independent mechanisms. First, the junctional permeability barrier promotes the formation of hydrostatic pressure via maintaining ion gradients which drive osmotic water influx. Second, tight junctions inhibit myosin contractility, which reduces apical tension and allows lumen inflation at lower hydrostatic pressure.

### Minimal force balance model recapitulates and predicts lumen phenotypes

To better understand how the changes in hydrostatic pressure and junction tension relate to 3D shape changes of lumen, we used an analytical model to describe the force balance of the luminal cavity (Fig. 5A, Material and Methods and Suppl. Material). We assumed that the volume and shape of the lumen are determined by the minimisation of free energy of the lumen surface that includes contributions from hydrostatic pressure and the junctional tension. In addition, we included the finding that the apical surface per cell does not adapt to changes in lumen volume in our experiments (Fig. 2). Therefore, we introduced a constant apical area per cell (experimental value). This surface also contributes to the force balance via a bending modulus (Suppl. Material). Thus, deforming the apical surface is energetically costly.

**Figure 5:**
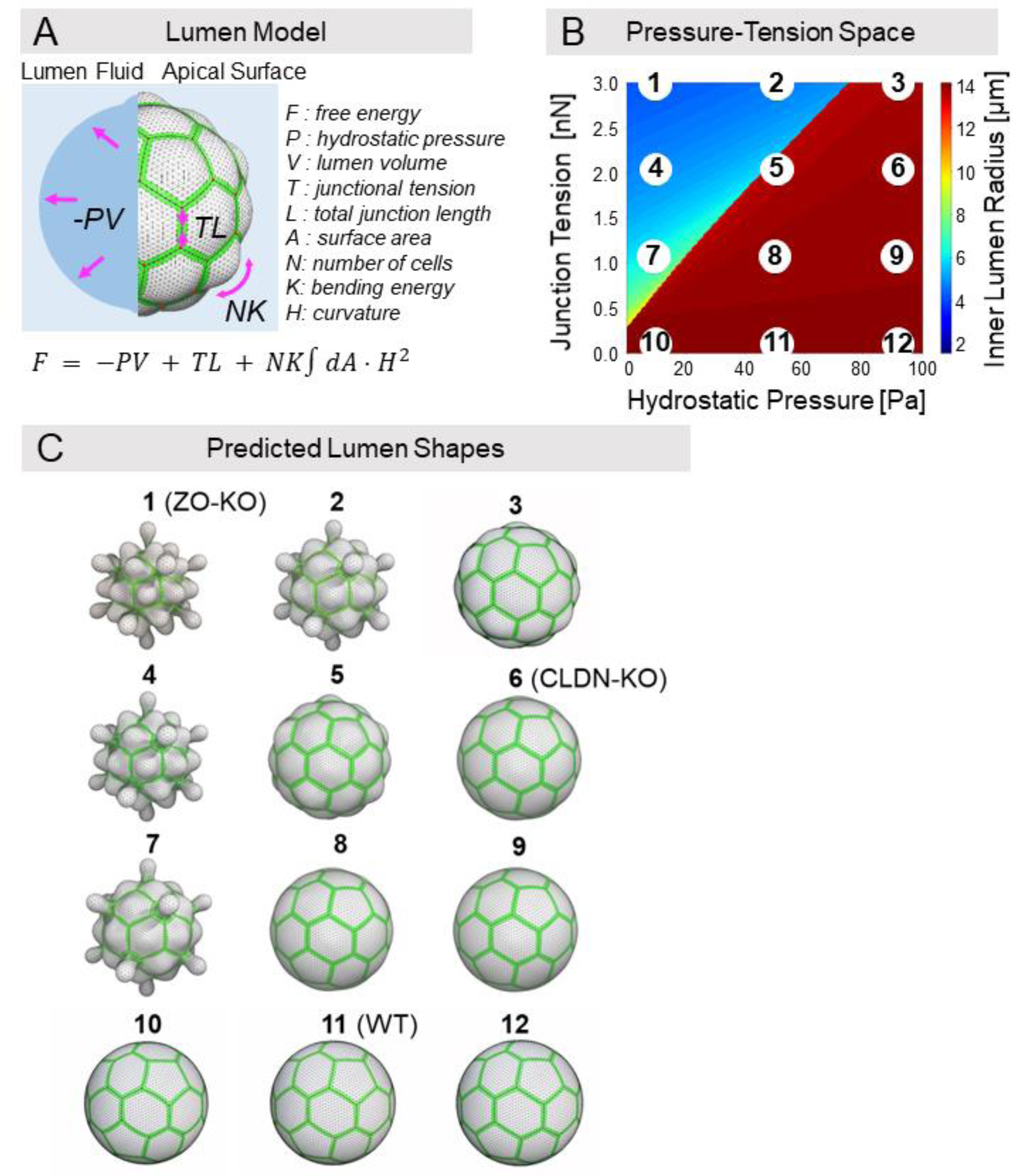
Apical surface model recapitulates lumen phenotypes of TJ-mutants. (A) Schematics of the free-energy model of lumen surface. The shape of the lumen is obtained by minimizing the free energy *F* of its area *A* computed with contributions from the hydrostatic pressure *P* of the lumen, the apical junctional tension *T* and the bending rigidity *K* of apical surface. (B) Phase diagram of the inner lumen radius in the apical line tension – hydrostatic pressure parameter space. For this phase diagram, the bending energy is *K* = 2. 10^-16^ J and the number of cells is N_cells_ = 42 cells. (C) Minimum energy 3D shapes of the lumen surface for the points labelled 1 to 12 in (B), calculated using Surface Evolver.

Varying lumen pressure and junctional tensions in the regime of experimentally determined values resulted in two lumen states (Fig. 5B). First, when junctional tension dominates the lumen volume is collapsed. Second, when hydrostatic pressure dominates the lumen volume is inflated to a spherical shape. Introduction of constant apical surface area with a bending modulus prevented full collapse of the lumen volume in the tension dominated regime (Suppl. Fig. 2A, B). Thus, the bending energy of the constant apical surface area promotes lumen opening especially in the regime when hydrostatic pressure is not sufficient to inflate the lumen (Suppl. Fig. 2C).

To compare the morphologies of the lumen predicted by our model with the experimental observed shapes, we calculated the energy minimized shape of a lumen surrounded by 42 cells (truncated icosahedron) for 12 positions in the pressure-tension range of our experiments with a vertrex model using Surface-Evolver^20^ (Fig. 5B). The calculated shapes revealed a striking qualitative agreement with the experimental observations (Fig. 5C). In the tension dominated collapsed state the excess apical membrane bends away from the lumen forming the characteristic lobes that are observed in the ZO-KO cysts (Fig. 5C states:1,2,4,7). In the fully inflated state, the apical surface is a smooth sphere resembling WT cysts (states: 8,9,6). Close to the transition line apical membranes slightly bend outwards (states: 5+3), which resembles CLDN-KO cyst morphologies.

In summary, force balance model is sufficient to recapitulate lumen morphologies of WT and TJ-mutants of MDCK cysts in the experimentally determined parameter regime. Thus, the model supports the notion that TJs control lumen morphology by tuning the mechanical force balance of tension vs pressure and bending of the apical surface. Mapping the pressure-tension parameter space, showed how TJ in WT tissue promote efficient inflation of spherical lumen at relatively low hydrostatic pressure by decreasing apical-junctional tension. The model also highlights the mechanical role of the apical membrane, which due to its conserved area and resistance to bending prevents full collapse of the lumen at high tensions and effectively facilitates lumen opening in the low tension vs pressure regime, which is in line with findings from the Dunn lab^5^.

To further test the mechanical model, we performed perturbations experiments focusing on the rescue of the collapsed lumen phenotype of ZO-KO cysts. To facilitate lumen opening we reduced junctional tension via myosin inhibition and we increased luminal pressure via micropipette inflation.

### Increasing hydrostatic lumen pressure partially reverses lumen collapse in ZO-KO cysts

To increase luminal pressure of MDCK cysts, we developed a micropipette inflation approach. For this purpose we grew cysts in shallow PDMS micro-cavities^21 22^, which allowed us to apply a fluid jet directly against the surface of cysts without detaching the cyst from the surface and the Matrigel. This setup enabled us to efficiently quantify morphological changes upon pipette inflation using confocal microscopy with high throughput (Fig. 6A). We found that the fluid jet applied to the cyst surface was able to open tight junctions and press additional fluid into the lumen increasing the hydrostatic lumen pressure. This was confirmed by seeing fluorescent dextran entering the lumen from the pipette solution (Fig. 6B).

**Figure 6:**
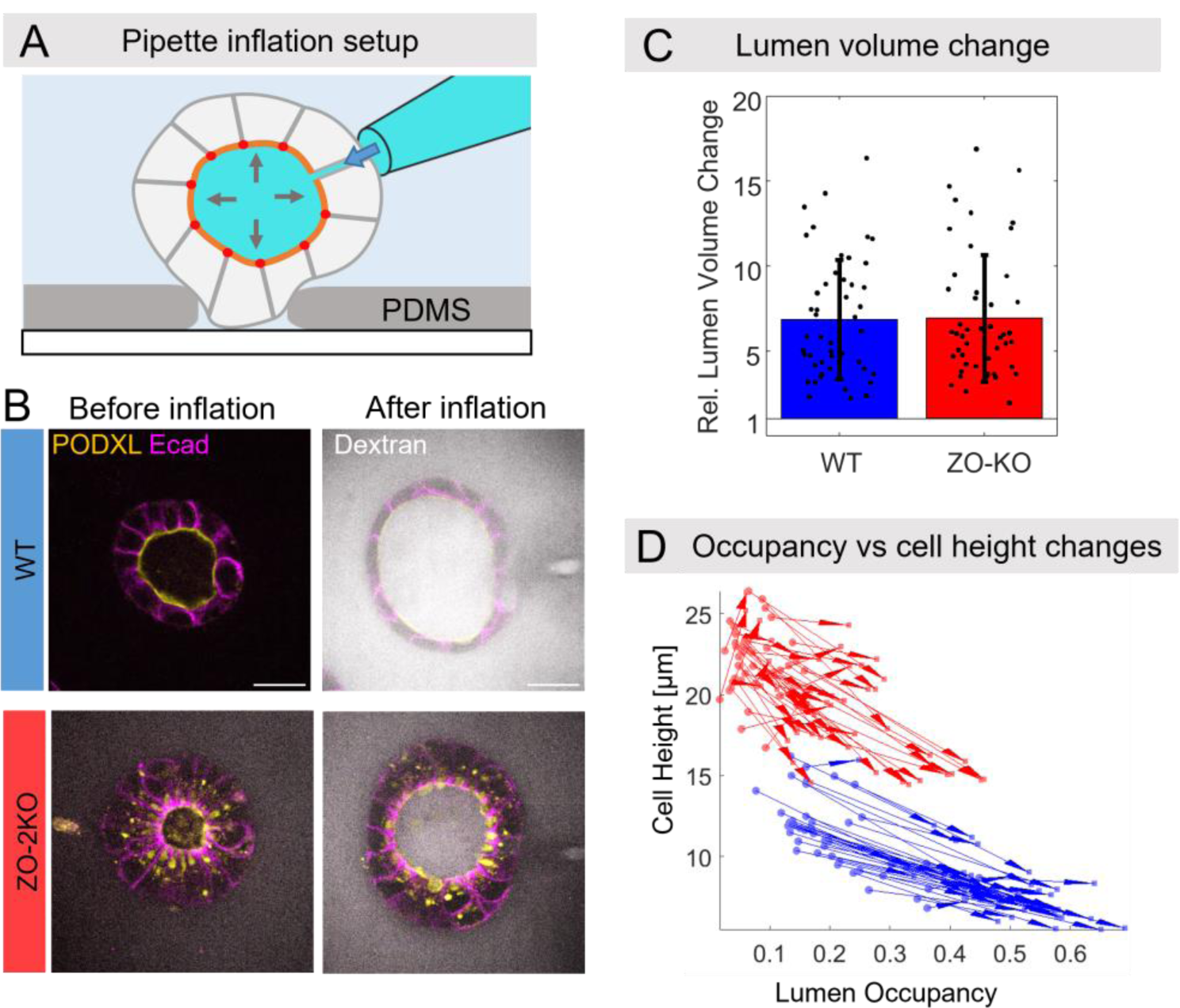
Increasing hydrostatic lumen pressure promotes lumen opening. (A) Schematic of pipette inflation assay. (B) WT and ZO-KO cysts before and after lumen inflation. Fluorescently labelled Dextran in the pipette solution was used to demonstrate entry of fluid into the lumen. (C) Quantification of lumen volume change normalized to the volume values before inflation. (D) Trajectories of lumen occupancy vs cell height before and after inflation represented by arrows show that ZO-KO cysts cannot be inflated to the same level as WT cysts and cells remain more elongated. WT in blue and ZO-KO in red. WT (n =96), ZO-KO (n=96) for C,D.

Fluid addition to the lumen resulted in significant expansion of WT and ZO-KO cysts. In both cases the lumen volume on average increased 7-fold (Fig. 6C). However, since ZO-KO cysts started with much smaller lumen, the absolute volume after inflation remained smaller compared to inflated WT cysts. To quantify how the cell monolayer responded to the lumen volume increase we quantified the change in cell height relative to the initial values (Fig. 6D). We found that the average cell height decreased 0.7 ± 0.1-fold in WT cyst and significantly less in ZO-KO with 0.9 ± 0.2-fold. Overall, ZO-KO cells started significantly more elongated compared to WT and even after lumen inflation did not reach the cell height of pre-inflation WT cysts. Relating the cell height change to the change of lumen occupancy after lumen inflation (Fig. 6D), showed that WT and ZO-KO cysts follow the same trend, e.g., increase in lumen occupancy and decrease in cell height. However, ZO-KO cysts could not be inflated to the same degree as WT cysts. While lumen occupancy in WT cysts increased from 0.2 ± 0.1 to 0.5 ± 0.1, the cell height decreased from 10 ± 2 µm to 8 ± 2 µm. In contrast, lumen occupancy of ZO-KO cysts only increased from 0.08 ± 0.04 to 0.22 ± 0.1, while the cell height remained relatively high with a change from 22 ± 2 µm to 19 ± 3 µm.

Taken together, increasing hydrostatic lumen pressure in WT and ZO-KO cysts via pipette inflation caused lumen expansion in both cases. However, cells in ZO-KO cysts remained significantly more elongated and lumen volume remained smaller compared to WT cysts. Thus, ZO-KO cells resisted lumen inflation and cell thinning in response to increased hydrostatic pressure significantly more than WT tissue, which is in line with the increased junction tension of ZO-KO tissue.

### Reduction of junctional tension increases lumen volume in ZO-KO and CLDN-KO cysts

To reduce junctional tension we inhibited Rho-kinase (ROCK), an upstream myosin IIa activator, via the compound Y-27632^23, 24^. Inhibition of ROCK via 200 µM Y-27632 resulted in efficient reduction of myosin accumulation at apical junctions after 120 min to almost WT-levels in ZO-KO and CLDN-KO monolayers (Suppl. Fig 3A,B). In line with the 2D monolayer experiments, Myosin-IIa was mostly enriched at the basal membrane in WT cysts, basal and apical-junctional in CLDN-KO cysts and basal, lateral and apical-junctional in ZO-KO cysts. Y-27632 treatment on 3D cysts resulted in qualitatively similar behavior as in 2D monolayers, e.g., myosin accumulation at the cell membrane including the apical junction complex was decreased (Fig. 7A, B). Since junctional myosin levels strongly correlate with junctional tension (Fig. 4F), Y-27632 treatment is expected to decrease junctional tension.

**Figure 7.**
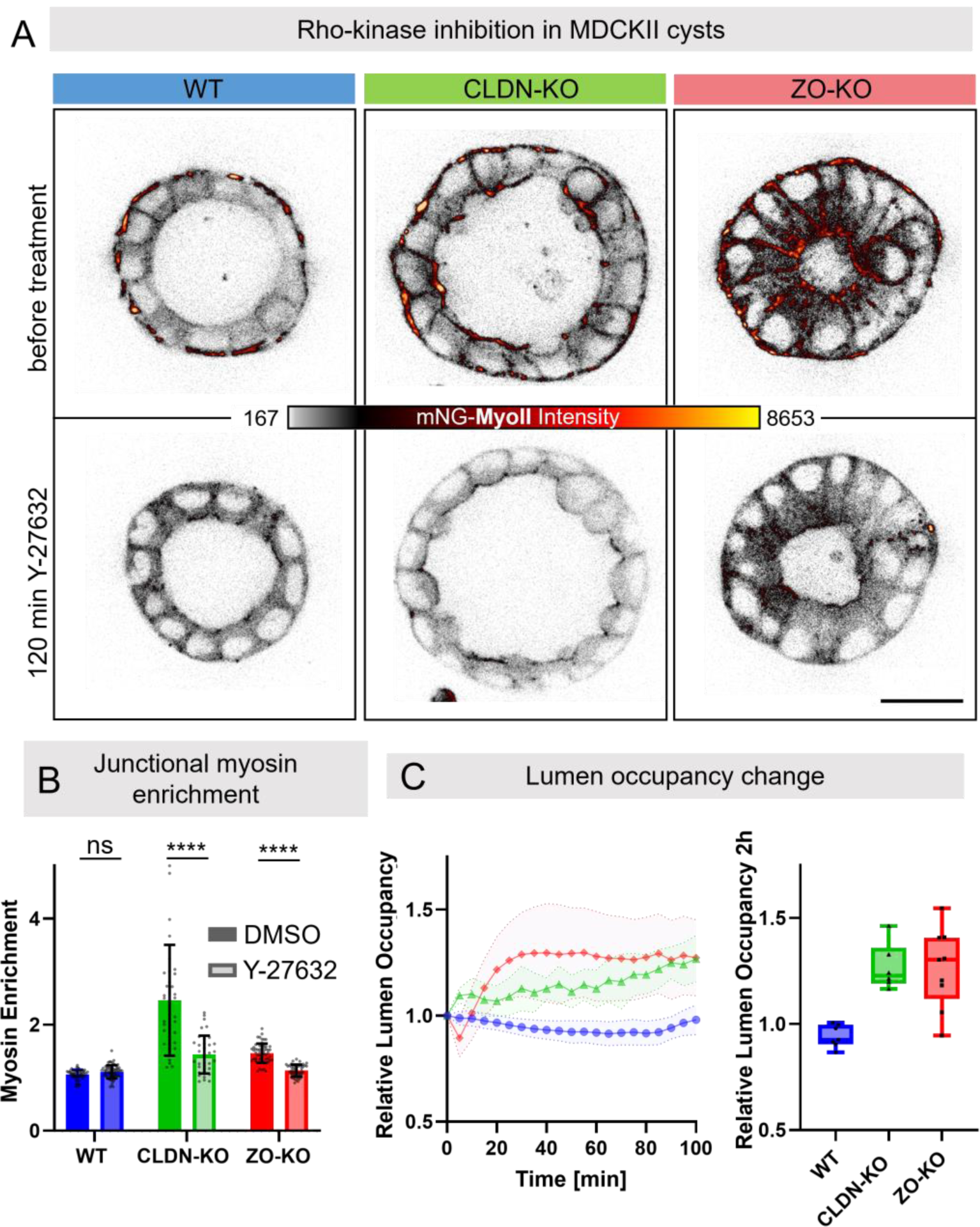
Release of junctional tension promotes lumen opening in CLDN-KO and ZO-KO cysts. (A) Distribution of endogenous myosin-IIa tagged with mNeon in MDCK-II cysts before and after addition of Rho kinase inhibitor Y-27632 (200μM) in WT and CLDN-KO and ZO-KO cysts. Scalebar 20µm. A clear increase in lumen size is visible in ZO-KO cysts. (B) Quantification of junctional myosin enrichment at cell-cell junctions versus cytoplasm and comparison between incubation with Y-27632 and the solvent DMSO 120 minutes after treatment, WT (n =57), WT Y-27632 (n=64), CLDN-KO (n=41) CLDN-KO Y-27632 (n=37), ZO-KO (n=57) and ZO-KO Y-27632 (n=) cysts (n=48).. (C) (Left) Normalized lumen occupancy after Y-27632 addition. WT cysts showed no significant change of lumen volume. ZO-KO cysts significantly increased lumen occupancy 30min after myosin inhibition. WT (n =7), CLDN-KO (n=6) and ZO-KO cysts (n=9). (Right) Mean change of lumen occupancy after 120min of Y-27632 addition. WT cysts show no significant change of lumen volume after tension release while CLDN-KO and ZO-KO cyst show a 40% increase of their lumen volume. WT (n =7), CLDN-KO (n=6) and ZO-KO cysts (n=9).

In WT-cysts, addition of Y-27632 had no significant influence on cyst and lumen morphology (Fig. 7C). In contrast, in ZO-KO cysts we observed a significant increase of lumen volume starting 20 min after Y-27632 addition and reaching a plateau after 40 min. Overall, the lumen volume of ZO-KO cysts increased 1.2-fold after inhibition of myosin (Fig. 7C). In CLDN-KO cysts, inhibition of myosin also resulted in an increase of lumen volume, however with a slower kinetic and to a lesser extent compared to ZO-KO cysts.

In summary, ROCK inhibition resulted in release of junctional myosin-IIa accumulation in TJ mutant tissues to almost WT levels. The release of myosin caused strong lumen expansion in ZO-KO and medium expansion in CLDN-KO cysts, while WT cysts were not affected. The perturbation results support the force balance model and show that releasing apical-junctional tension is an important function of TJs to promote lumen inflation by hydrostatic pressure.

## Discussion

In this work, we addressed the question how tight junctions contribute to formation of lumen in an organotypic 3D tissue culture model system (MDCKII cysts). By measuring and modeling the force balance of hydrostatic lumen pressure and junctional tension of TJ mutants, we found that TJs have a strong impact on the amount of apical-junctional tension and to a lesser degree on the amount of hydrostatic pressure established by solute pumping. In addition, our measurements revealed that cells control the area of the apical membrane independently from hydrostatic pressure, junctional tension and lumen volume.

### Role of TJs in regulating tissue permeability and hydrostatic lumen pressure

Comparing hydrostatic lumen pressure of wild-type and TJ-mutant cysts revealed that the paracellular barrier for ions formed by Claudins strands is not required to build up hydrostatic lumen pressure and inflate spherical lumen (CLDN-KO, Fig. 2, 3). Previous trans-epithelial permeability measurements of small molecule tracers showed that permeability for molecules with a size of ≈ 1 nm is ≈ 9-fold increased for CLDN-KO tissue and ≈ 12-fold increased in ZO-KO tissue compared to WT^7^. In addition, ZO-KO tissue is also permeable for larger macro-molecules of up to ≈ 15 nm ^7^. The significantly increased trans-epithelial leak for solutes of CLDN-KO and the ZO-KO tissue should, in principle, reduce the osmotic pressure difference between the lumen because the para-cellular leak channels equilibrate concentration gradients established by active solute pumping into the lumen^16^. Indeed, previous work focusing on pressure independent mechanisms of lumen formation assumed that, due to the increased solute permeability, CLDN-KO cysts have reduced hydrostatic pressure^5^. However, we found that, despite the strongly increased permeability, CLDN-KO cysts establish a hydrostatic lumen pressure comparable to WT tissue and are able to inflate spherical lumen (Fig. 2, 3). This suggests that cells can compensate increased para-cellular solute permeability. One mechanism could be that cells increase active ion pumping into the lumen to compensate the increased para-cellular leak. On the other hand, our results suggest that ZO-KO cysts are not able to establish significant hydrostatic pressure (Fig. 3). This could imply that cells are not able to compensate the further increased solute permeability in the ZO-KO which would reveal an upper limit for the pumping capacity. However, since the ZO-KO cysts also have strongly increased junctional tension and very convoluted lumen shapes it is possible that our lumen volume analysis was not able to resolve small changes in volume after drainage in particular of the folded lobes of the lumen. Thus, it remains possible that also ZO-KO cysts can maintain an osmotic pressure in the lumen. Along these lines, it has been suggested that MDCKII cysts grow in a regime in which the osmotic pressure ΔΠ is significantly larger than the hydrostatic pressure ΔP (ΔΠ ≫ ΔP)^16^. Thus, a significant reduction of osmotic pressure due to increased paracellular leaks may still lead to conditions in which lumen can expand by osmotic water influx (ΔΠ > ΔP) albeit at slower speeds. Further development of methods that can directly measure local osmotic pressure will be required to understand the feedback between paracellular permeability, solute pumping and hydrostatic pressure.

Overall, our results suggest that epithelial cells can maintain sufficient osmotic lumen pressure to drive water influx and promote lumen expansion even when tight junctions open the para-cellular barriers. This points to a feedback mechanism that regulates solute pumping activity to maintain a certain luminal pressure. Recent work suggests that MDCK cells can sense the apical-basal pressure difference and modulate active fluid pumping via localization of the Na/K ATPase^18^.

### Role of TJs in regulating apical-junctional tension during lumen formation

The TJ scaffold proteins ZO1 and ZO2 have been shown to be important regulators of the apical actin cortex^9–11^ . In addition, ZO1 is known to inhibit myosin activity via inhibition of the RhoA exchange factor GEF-H1^25^, and depletion of ZO1 leads to increased junctional tension^26, 27^. Our quantification of endogenous junctional myosin-IIa levels in combination with junction tension measurements confirm the key role of ZO proteins in regulating apical-junctional contractility via myosin inhibition (Fig. 4,7, Suppl. Fig. 3). In addition, we found that claudins also play a critical role in tension regulation. Depletion of claudins causes an increase of myosin and junctional tension, albeit to lesser extent than ZO1 depletion. Since junctional ZO1 levels are not affected by claudin depletion, myosin regulation via claudins likely involves a different pathway. Along these lines, recent work on transient tight junction breaching via mechanical stretching has revealed that breaks in the tight junction network cause local calcium bursts, which activate RhoA and lead to a contractile response via myosin^28, 29^. This response leads to efficient healing of TJ breaks and ensures active maintenance of the TJ barrier in a mechanically challenging tissue environment. While the mechano-sensor that responds to TJ stretching has not been identified, we speculate that the same mechanical feedback may also be activated in ZO-KO and CLDN-KO tissue, which could explain the increase in junctional tension of CLDN-KO cells despite normal ZO1 levels.

Taken together, our results are in line with previous reports that TJ regulate junctional tension via control of acto-myosin activity^9–11^ . Importantly, we found that a key role of TJ for lumen formation is to decrease apical-junctional tension which promotes spherical lumen inflation at lower hydrostatic pressures.

### Tuning of lumen shapes by TJs and the role of the apical membrane cortex

In line with previous observations^5^, we found that the apical surface area per is cell is mostly constant during lumen growth (Figure 2). Even when the lumen volume is collapsed in ZO-KO cyst the apical area remains close to that of WT cells. This suggests that cells control the size of the apical membrane independently of hydrostatic pressure and tension probably by tuning the balance of apical endo- and exocytosis^30, 31^ and/or by independently regulating the apical actin cortex. This may also suggest that the size of the apical area is genetically encoded in a cell type specific manner rather than being a parameter that can freely adapt to morphological and mechanical changes of cells and tissues. Therefore, we included an apical area constraint in our mechanical force-balance model of lumen morphology (Fig. 5). Our model is able to capture the key morphological changes of CLDN-KO and ZO-KO compared to WT cysts and therefore provides a useful tool to understand morphological transitions during lumen growth and remodeling. In future work, it will be important to (i) further consolidate the role of the mechanical properties of the apical membrane and its area regulation, (ii) include the contribution of lateral and basal surface mechanics, and (iii) include the role of active pumping and osmotic pressure into the modelling framework to explore the dynamics of lumen formation^3, 16, 32^ .

## Supporting information

Suppl_Methods_Model

## Acknowledgements

We thank Mikio Furuse, National Institute for Physiological Sciences (NIPS) Japan, for providing the CLDN-5KO MDCKII cell line. D.R. team thanks support from the Imaging Platform of IGBMC. DR team was supported by the Interdisciplinary Thematic Institute IMCBio, as part of the ITI 2021-2028 program of the University of Strasbourg, CNRS and Inserm, was supported by IdEx Unistra (ANR-10-IDEX-0002), and by SFRI-STRAT’US project (ANR 20-SFRI-0012) and EUR IMCBio (ANR-17-EURE-0023) under the framework of the French Investments for the Future Program. AH and M.M. were supported by the Volkswagen Foundation project number A133289. M.J.B and M.F.S were funded by the Max Planck Society, including through an ELBE fellowship. This project was funded by HFSP project number RGP0050/2018.

## Author Contributions

A.H. conceived the project. D.R. and T.G. conceived cyst drainage assay for the estimate of luminal pressure, A.H. developed the experimental implementation and M.M., R.M. performed measurements. D.R. conceived the pipette assay with microfabricated substrate and the lumen inflation with T.G. and L.L. who performed the experiments and measurements. C.H.W. performed morphological characterization of TJ-mutants. M.M. performed junctional tension measurements and myosin perturbation experiments. T.H., M.S., D.R. and T.G. conceived the mechanical model, T.G. performed the associated numerical simulations. M.J.B. and M.F.S. implemented the model in Surface Evolver. C.M-L. did the cell-culture work and created stable cell lines. A.H. and M.M. wrote the manuscript with input from all authors.

## Star Methods

### Cell culture maintenance and cell lines

MDCK-II cells were cultured in MEM (Sigma, USA), 5% FBS (Sigma, USA) with 1 mM of sodium pyruvate (Thermo Fisher Scientific, USA) and 1% of 100x MEM Non-Essential Amino Acids (Thermo Fisher Scientific, USA) at 37°C with 5% CO_2_. ZO1/2 knockdown MDCKII as well as MDCKII WT-ZO1-mNeonGreen were created by Beutel et al. using Crispr/Cas^13^ . Claudin quin-KO MDCKII cells were obtained from the Furuse lab^7^. For N-terminal endogenous labelling of myosin with mNeon in WT, ZO-KO, CLDN-KO cells, the myosin-2-A exon was targeted using Crispr/Cas.

### Three-dimensional culture of MDCK cysts and drug treatment

MDCK cysts were cultured as previously described^33^, in brief: MDCK monolayers were dissociated and either embedded in Matrigel (Corning, USA) for the pseudo-time-laps assay or seeded on 0.5 mg/ml laminin (Roche Diagnostics, Switzerland) coated 35-mm glass-bottom dish (MatTek Corp., USA) containing 5% of Matrigel for the other assays referred to in this manuscript. Cells were then cultured for 4-5 days and subsequently imaged. For perturbation of actomyosin contractility by ROCK inhibition, Y-27632 dissolved in DMSO was applied to the culture media with 200μM working concentration.

### Trans-epithelial permeability assay

3D adherent cysts were cultured as already described for 4 to 5 days in 37 °C 5% CO_2_ until reaching 30 to 40 μm in diameter. To measure permeability of small solute (Dextran-Alexa488) and a lipid tracer (DPPE-TMR), cyst medium was supplemented with 1μM DPPE-TMR, which was complexed with fat-free BSA in 1:1 molar ratio, for 8min at 37°C. After washing, the medium was supplemented with 10μM Dextran-Alexa488. After 15min incubation, cysts were imaged on a confocal microscope to evaluate distribution of the lipid tracer and Dextran-Alexa488.

### Lumen sphericity measurement

To determine how close the lumen shape resembles a sphere, the sphericity (Ψ) was calculated by comparing the surface area (A) and to the surface area of a perfect sphere with the measure lumen volume 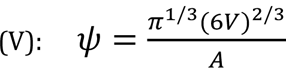. A sphericity of 1 refers to a perfect sphere while highly irregular lumen approaches a sphericity of 0.

### Drainage of lumen and quantification of hydrostatic lumen pressure

WT, CLDN-KO and ZO-KO cells stably expressing HaloTag-CAAX were cultured with the laminin-based method for 5 days to form cysts. Prior to the experiment, cysts were incubated in culture media with 200 nM of Janelia Fluor® 646 HaloTag® ligand overnight to visualize the membranes.

The cysts were cut open by ablating one cell using a Zeiss LSM 780 NLO system (Zen Black software version 11.00) with a 2-photon laser (Titatnium/Saphire) at 800 nm (Coherent, Santa Clara, USA) with a power of 3.2 W at the laser head and 100% laser output set in Zen Black. A line scan with a width of 3 µm and 5-line repetitions across the epithelium at the middle plane of a cyst was typically sufficient to open the lumen to the outside. Lumen collapse was imaged by taking 3D stacks of the cysts with a time resolution of 15 seconds.

To characterize the experiments quantitatively, we segmented the lumen in 3D over time and plotted the change of lumen volume as a function of time. These measurements allow us to calculate the initial flow as a key readout to evaluate hydrostatic luminal pressure. We also measured the channels dimensions using the passage of the fluorescent mucin (see Fig. 3C.). To obtain the hydrostatic lumen pressure before the cut, we modelled the flow going out of the cyst as a Hagen-Poiseuille flow of a liquid of viscosity η at a flow rate Q in a channel of radius R and length L:

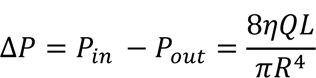

The viscosity of the lumen fluid η, was determined by measuring the diffusion of Dendra-mucin in the lumen using Fluorescence Correlation Spectroscopy experiments

### Measurement of lumen fluid viscosity

Viscosity of the luminal fluid was measured using fluorescence correlation spectroscopy (FCS) of dendra2-tagged soluble sialomucin (Dendra2-CD164) on an Abberior STED microscope (Abberior Instruments, Göttingen, Germany). The concentration of dendra2-mucin in the luminal fluid was typically too high for FCS. Therefore, we photo-converted a small fraction of dendra2 into the red-emitting form using a short scan of 405 nm light. FCS measurements were then performed using 561 nm excitation (20µW back focal plane) using a 60x water objective to measure the diffusion of the photo-converted dendra2-mucin in the lumen. FCS curves were fitted using a 3D diffusion model including triplet blinking as described previously^34^. To determine the viscosity in the lumen the diffusion times of dendra2-mucin in the lumen were compared to diffusion times of the same molecule in standard 2D cell culture medium with µ (DMEM @ 37°C) = 0.9 mPa*s (MDCK monolayer on glass) (Figure S1).

### Three-dimensional segmentation of MDCK cysts

Fluorescent Z-stacks of MDCK cysts were acquired by confocal microscopy. The images were processed with median filter to better preserve the structures. All reconstructions of three-dimensional images were performed with LimeSeg ^35^ plugin from Fiji185. The segmentation on the 4-D recordings of lumen formation was performed using a groovy script that automatically operated LimeSeg to multiple time points where segmented dots of present time point were applied to next time point as ROI for segmentation. The geometrical information measured with segmented mesh was automatically saved. The number of cells of a cyst was quantified with TrackMate of Fiji on the nuclei signals.

### Tension measurement

Laser ablation was performed on Zeiss LSM 780 NLO system driven by Zen Black software version 11.00. Images pixel size was 0.268 µm x 0.2268 µm. Objective used was Zeiss C-Apochromat 40x/1.2 W. Ablation was performed with 800 nm Titanium/Sapphire femtosecond pulsed laser Chameleon from Coherent (Santa Clara, USA) with a power of 3.2 W at the laser head, 60% laser output set in Zen Black, reflected by MBS 760+, with pixel dwell time for photomanipulation of 7.2 µs, single iteration, ablation area was line scan, 10 pixels. For measuring the recoil velocity, the lateral junction of MDCK WT or CLDN-KO and ZO-KO cells was highlighted by ZO1-mNeonGreen or MyosinIIa-mNeon green respectively with the following settings: mNeonGreen was excited with 488 nm (Argon Laser) with MBS 488/561/633, emission filter used was 490-570 nm, pixel dwell time 2.83 µs, approximately 7.7 fps with GaAsP detector. Recoil velocities were determined by dividing the distance covered by the vertices in a considered span of time, by the same time span.

### Immunostainings of MDCK tissue

Samples were fixed with 4% w/v Paraformaldehyde (PFA) pH 7.25 in PBS for 10 minutes at room temperature. For pseudo time-lapse imaging samples were fixed with4% PFA + 0.025% v/v Glutaraldehyde in PBS for 10 minutes at room temperature to prevent cells being washed away due to PFA incubation. Cells were then permeabilized with 0.5% v/v Triton X-100 in PBS for 10 minutes and blocked with the blocking buffer (2% w/v BSA, 0.02% v/v NaN3) in PBS for 1 hour at room temperature. Primary antibodies were diluted by the blocking buffer and incubated with cells for 1 hour at room temperature or overnight at 4°C followed by diluted secondary antibodies and labelled phalloidin incubation. Primary antibodies used were: Rabbit anti-gp135/podocalyxin (1:200; from Ünal Coskun’s group); Mouse anti-E–Cadherin (1:100; from MPI-CBG antibody facility).

### Confocal microscopy

Live cell imaging: the images were acquired with a Zeiss Axio Observer.Z1 inverted microscope (ZEISS, Germany) equipped with a CSU-X1 spinning disc confocal scanning unit (Yokogawa, Japan), driving a Zeiss AxioCam MRm, Monochrome CCD camera. The images were acquired with a x63 1.2 NA water objective lens (ZEISS, Japan) and a x100 1.46NA alpha Plan-Apochromat oil objective lens (ZEISS, Japan) and ZEN blue software (ZEISS, Germany). Z-stacks were acquired with a distance of 300 nm.

### Cyst inflation in microfabricated cavities

To increase the volume of the lumen, we applied a local liquid flow using a micro-pipette to cysts growing in microfabricated cavities, which prevented detachment and movements of cysts (see below). Application of liquid flow led to the rapid inflation of the cyst within seconds, which was reversible (see Fig. 6). In addition, we found that fluorescent Dextran, present in the pipette solution, entered the lumen, which indicates that the liquid stream opened cell junctions and presses liquid into the lumen (see Fig. 6A-B.).

The inflation experiments were quantified by segmenting lumen and cyst shape before and after inflation to extract tissue and cell shape changes in response to the hydrostatic pressure perturbations.

The micro-cavities were prepared by soft lithography^21^. Briefly, a mask with 18 μm disks with 100 μm inter-distance was designed with Autocad, which was then printed on photomask. The pattern was transferred to SU8 silicon wafer by photolithography. Then a PMDS mold was generated by soft lithography. The PDMS pillar replica were transferred on a cover-glass using plasma activation and chlorotrimethylsilane for passivation and subsequently filled with liquid PDMS. With overnight polymerization at 65°C, the PMDS formed a solid membrane on the cover-glass, and cavities could be obtained by gentle removal of the PDMS stamp. After obtaining through-hole cavities on the cover-glass, the surface was activated by plasma and then incubated with 5 µg/ml laminin at room temperature. Cells were seeded in cavities by centrifugation^36^. Then 15µl Matrigel was added on the top of the cavities, which allowed MDCK to form cysts. Cysts were used 5 days after cell seeding for inflation experiments. To facilitate the experiments, Matrigel on top of micro-wells was removed using a needle.

The lumen was inflated by using a pipette pulled with a Sutter pipette puller (P 2000), the tip of the pipette was then broken using a scalpel under a binocular to the expected diameter. The flow was controlled with the Eppendorf Cell Tram Vario. We injected medium (MEM with 5% serum) containing fluorescent Dextran to track the localisation of the liquid. Finally, changes in cell and cyst shapes were monitored live by imaging by using Leica spinning disk (Spinning disk LEICA CSU W1).

### Lumen Model

The lumen shape was calculated based on the minimization of the free energy given by the sum of the contributions from the hydrostatic pressure in the lumen, the line tension at apical junctions of all neighbouring cell pairs and apical bending energy from the lumen surface.

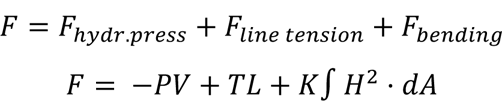

where *P* is the pressure of the lumen, *V* its volume, *T* the apical line tension, *L* the total apical line length, *K* the effective bending energy of the apical membrane, *H* its mean curvature. To obtain an analytical solution to this problem, several geometrical assumptions must be made which are detailed in the supplementary note 1.

To estimate the junctional line tension *T* = ζ*v* from the recoil velocities v determined by laser ablation experiments (Figure 4) we used a damping coefficient of ζ = ɳ_*cortex*_ℓ_*cortex*_ = 50*pN.s/εm*. For this we take the typical value of η_cortex = 200 Pa. s ^37^ and a cortical length of l_cortex= 500nm. Using these values with v = 10 µm/s gives line tensions in the range of T = 1000 pN (see parameter table below), which corresponds to about 10^4^ myosin motors applying a 0.1pN mean force^38^ . We assume the same ζ damping coefficient for the different MDCKII-mutants because the change in myosin expression levels is related to the changes in tension and not in viscosity of the cortex.

Values for the cortex modulus have been previously measured to be 2·10^2^k_B_T^39^ . To take the rigidity of the cell layer into account, we estimated an effective bending modulus of 2·10^5^ k_B_T = k = 2.10^-16^ J (see Table 1). We integrate in this value the effective resistance of the cell layer by assuming a typical height of about 10 µm and a rough estimate for the associated change in bending modulus following similar scaling arguments proposed in^40^ . 3D shapes of lumen were calculated using Surface Evolver^20^ by minimizing the energy:

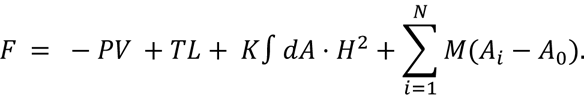

By term, the contributions are: the contribution from the fluid inside the lumen, where *P* is the pressure and *V* is the volume, contribution from the junctions, where *T* is the tension and *L* is the total junction length, bending of the apical surface, where *K* is the bending modulus and *H* is the mean curvature which we integrate over the surface, and finally area conservation, where *M* is the area modulus, *A_i_* is the area of cell *i* and *A_0_* is the preferred cell area (equal to *S*/*N*, where *S* is the total surface area and *N* is the total number of cells). Energy minimization was accomplished using the Surface Evolver programming language ^20^. Briefly, a surface consisting of 42 polygonal faces was imported into Surface Evolver. These 42 faces consist of a mixture of pentagons and hexagons, as dictated by Euler’s formula. The edges between the polygonal faces were marked to represent junctions. The surface was then triangulated and transformed into a sphere of surface area *S*. The energy function above was then applied and the surface evolved toward minimum energy with Surface Evolver’s gradient descent tools. A high area modulus was used to ensure cell areas remain approximately constant.

Parameters taken for computation of the phase diagram of Fig.5:

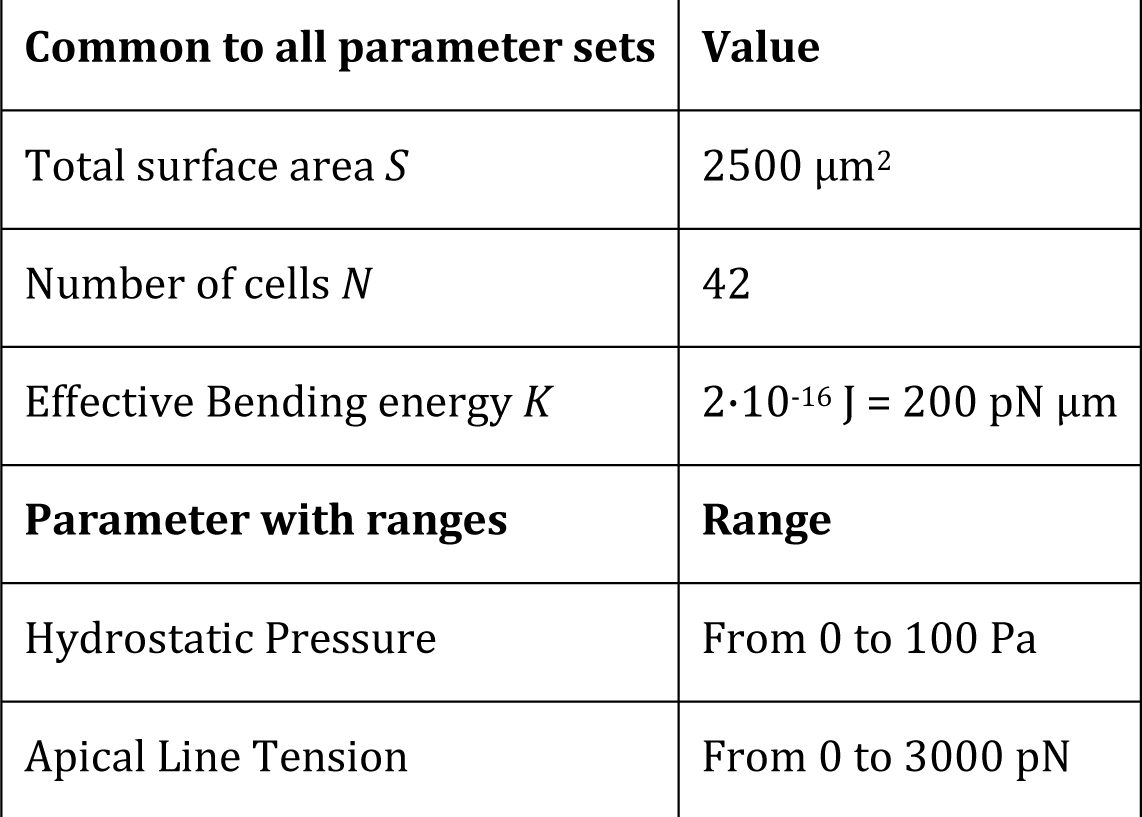

Parameters taken for computation of the lumen shapes (surface evolver) of Fig.5:

| Common to all parameter sets | Values |
| --- | --- |
| Total surface area $S$ | 2500 $\mu\text{m}^2$ |
| Number of cells $N$ | 42 |
| Bending energy $K$ | $2 \cdot 10^{-16} \text{ J} = 200 \text{ pN } \mu\text{m}$ |
| Area modulus $M$ | $1 \cdot 10^{-14} \text{ J}/\mu\text{m}^2 = 10000 \text{ pN } / \mu\text{m}$ |

| Parameter set label | Tension $T$ (pN) | Pressure $P$ (Pa = pN/micron<sup>2</sup>) |
| --- | --- | --- |
| 1 | 3000 | 10 |
| 2 | 3000 | 50 |
| 3 | 3000 | 90 |
| 4 | 2000 | 10 |
| 5 | 2000 | 50 |
| 6 | 2000 | 90 |
| 7 | 1000 | 10 |
| 8 | 1000 | 50 |
| 9 | 1000 | 90 |
| 10 | 0 | 10 |
| 11 | 0 | 50 |
| 12 | 0 | 90 |

---

**Suppl. Figure 1:**
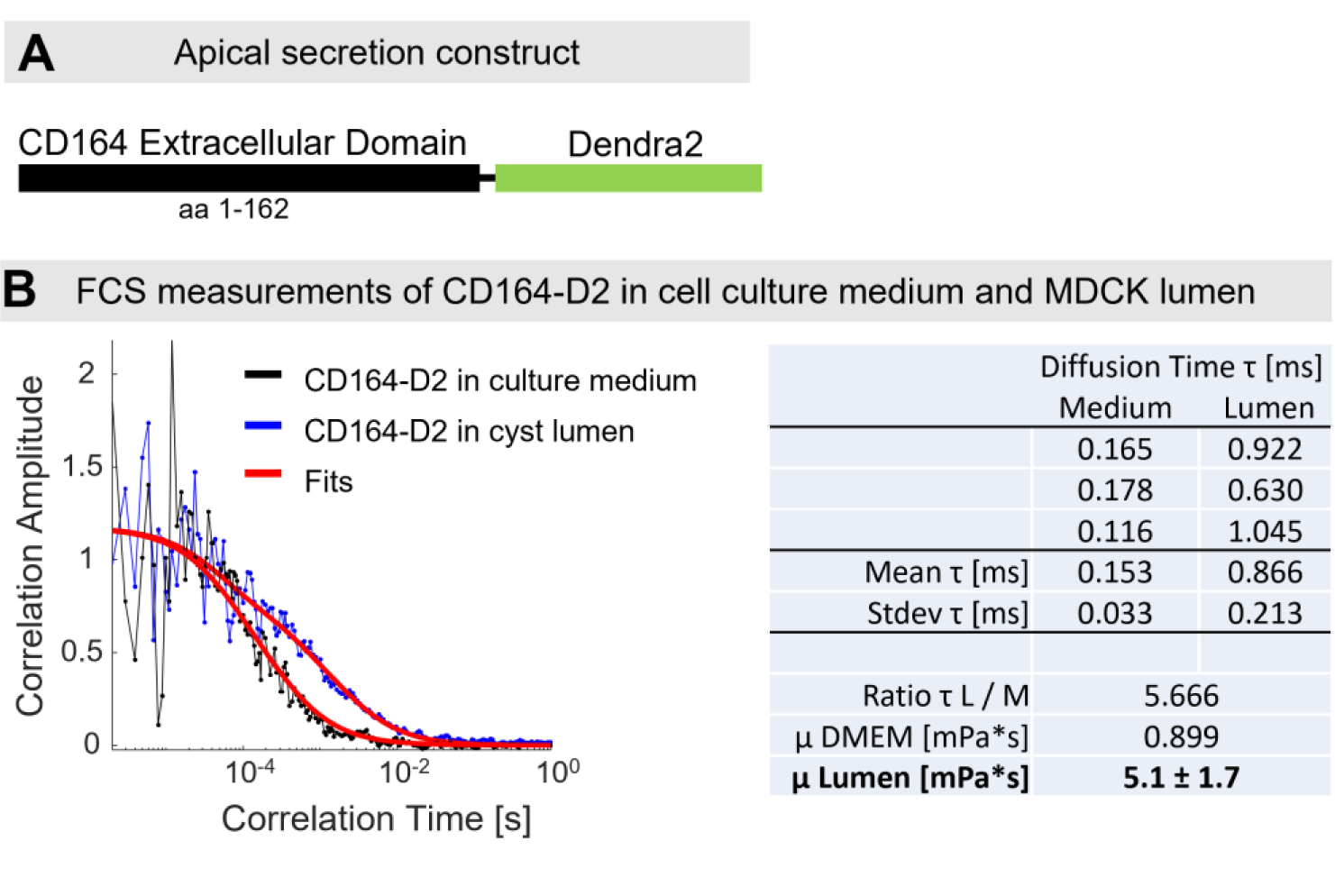
Measurement of lumen fluid viscosity. (A) Schematic of apically secreted CD164 construct. (B) Fluorescence correlation spectroscopy measurement was performed at 37°C to compare diffusion times of Dendra2-CD164 in the lumen and in cell culture medium. Viscosity in the lumen was determined to be η = 5.1 ± 1.7 mPa*s.

**Suppl. Fig. 2:**
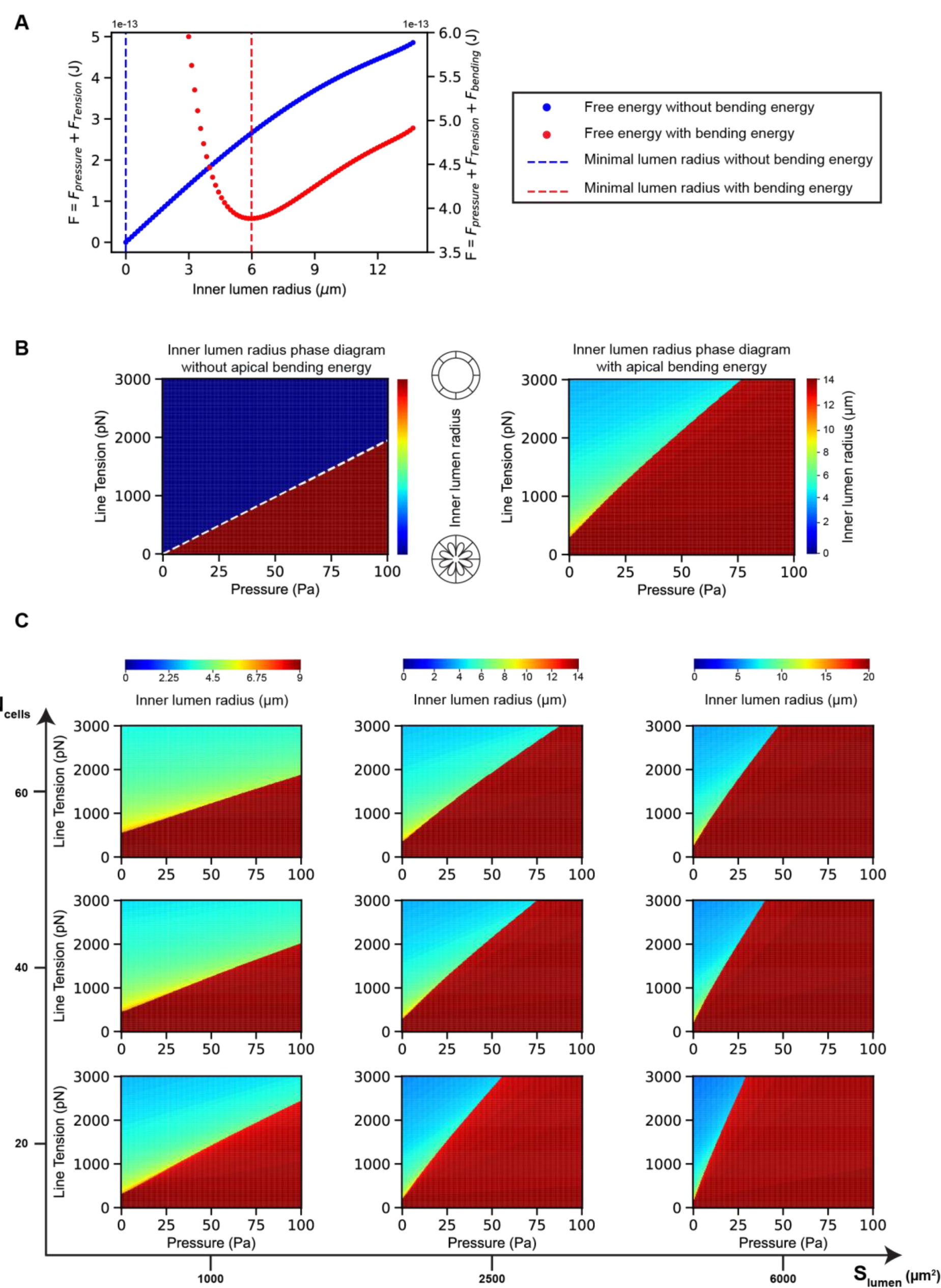
Contribution of bending energy, cell number and surface area on phase diagrams of lumen morphology. (A) Free energy of the lumen surface as a function of inner lumen radius computed for a total number of cells-N_cells_ = 42 cells, a total lumen surface - S_lumen_ = 2500 µm^2^, hydrostatic pressure of lumen - P = 20 Pa, apical junctional tension - T = 1500 pN. The minimum of free energy determines the value of the inner lumen radius at equilibrium. Blue curve represents the case without apical membrane bending energy, the minimal inner radius in this case is R = 0 µm. Red curve represents the case accounting for an apical bending energy (K = 2.10^-16^ J), the minimal inner radius in this case is non-zero. (B) Comparison of the phase diagrams of the inner radius of lumen determined by minimizing the free energy of the lumen surface without (left panel) and with (right panel) bending energy of the apical surface. Blue zones correspond to flower-like lumen shape and red zones to spherical inflated lumen. Total number of cells - N_cells_ = 42 cells, total surface of lumen - S_lumen_ = 2500 µm^2^, bending energy K = 2.10^-16^ J. (C) Phase diagrams generated for number of cells N_cells_ – total surface of lumen S_lumen_ pair values for a bending energy - K = 2. 10^-16^ J. Note that the inflated region (red zone) increases with larger S_lumen_ values and decreases with larger N_cells_.

**Suppl. Fig. 3:**
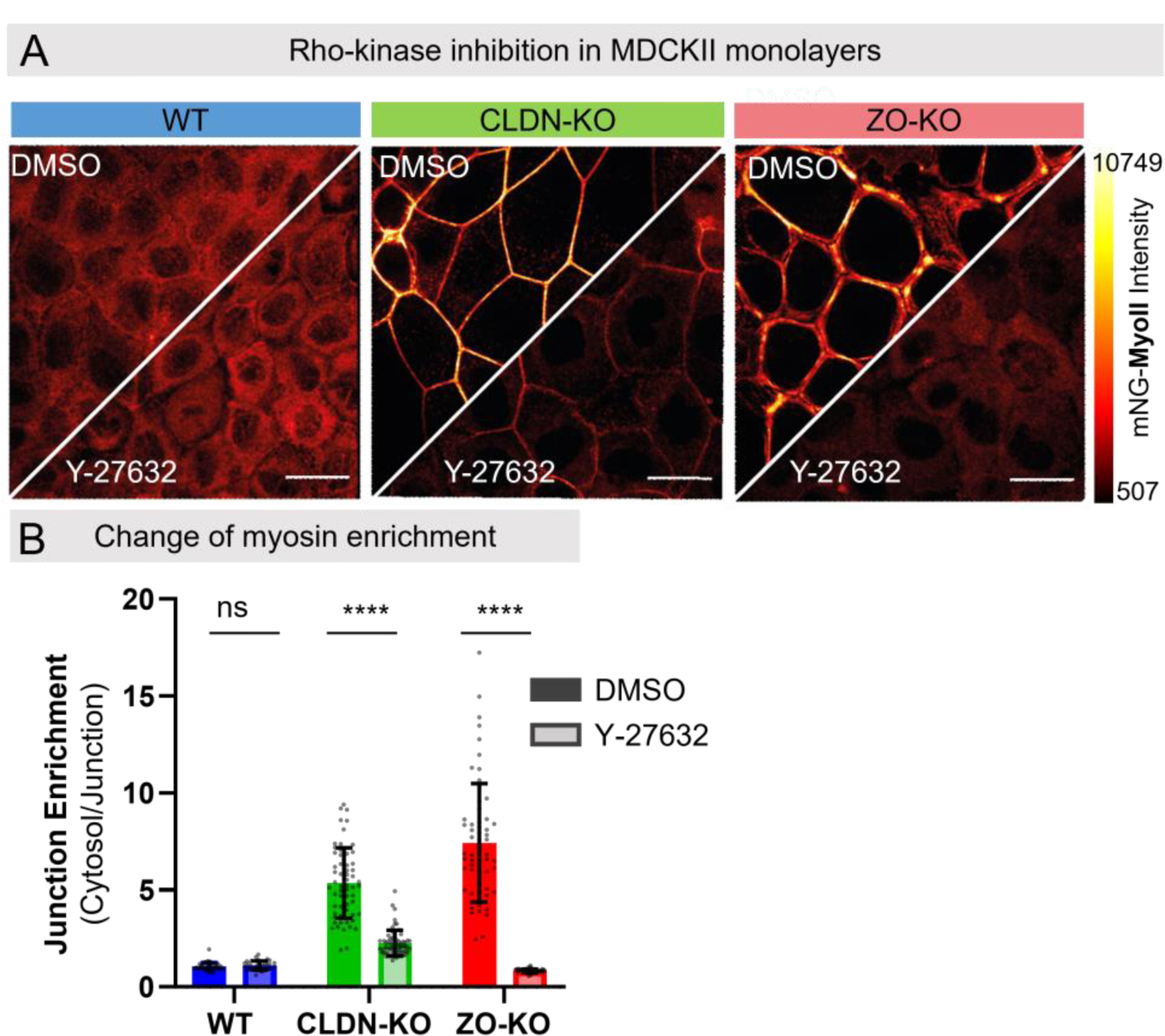
Release of junctional tension by ROCK inhibition. (A) Distribution of endogenous myosin-IIa tagged with mNeon in MDCK-II monolayers before and 2h post Y-27632 treatment (200µM). Scalebar 5µm. (B) Quantification of release of junctional myosin enrichment after Y-27632 with DMSO incubation as control . WT (n =50), WT Y-27632 (n=35), CLDN-KO (n=61) CLDN-KO Y-27632 (n=66), ZO-KO (n=60) and ZO-KO Y-27632 (n=) cysts (n=64).

