## Supplementary material for "Tight junctions regulate lumen morphology via hydrostatic pressure and junctional tension": Suppl_Methods_Model

### Supplementary Information: Analytical theory of lumen surface morphology

#### Contents

**1 Description of the problem** **1**

**2 First case: apical surfaces with no bending energy** **3**

**3 Second case: apical surfaces with a bending energy** **4**

**4 Differences between the two phase diagrams** **5**

#### 1 Description of the problem

We consider a spheroid composed of $N$ MDCK cells surrounding a lumen filled with fluid, the spheroid is embedded in an extracellular matrix (ECM) in three dimensions. The purpose of this supplementary material is to explain the observed morphologies that can be found in spheroids composed either of MDCK *wild type* cells or MDCK *double knockout ZO1-ZO2* cells (ZO-KO) (see Supp. Info. Fig. 1) through the analytical study of lumen-surface force-balance model.

We set lumen surface area or the total apical surface area to be a constant $S$ for a given number of cells $N$ in a spheroid. We assume that the cells are organised approximately in a hexagonal tesselation (Supp. Info. Fig. 2a). For each cell, the line perimeter of this hexagon is given by $P_{\text{hex}} = \sqrt{8\sqrt{3}S_{\text{hex}}}$. Next we assume that the surface of this hexagon is close to the surface of the circle it is inscribed in : $S_{\text{hex}} = \pi r_a^2$ with $r_a$ the radius of this circle (Supp. Info. Fig. 2b). Then the total line perimeter $L$ for $N$ cells is

$$L = \frac{1}{2}N\sqrt{8\sqrt{3}\pi r_a^2} = N r_a\sqrt{\pi\sqrt{6}}.$$

We assume each cell has an apical surface that can be described as a paraboloid with a base circle of radius $r_a$ and of depth $h$. Note that $h$ can be positive or negative. The volume of such paraboloid is $V_{\text{paraboloid}} = \frac{1}{2}\pi r_a^2 h$ and its surface is [1]

$$S_{\text{paraboloid}} = \frac{\pi r_a}{6h^2}\left[(r_a^2 + 4h^2)^{\frac{3}{2}} - r_a^3\right]$$

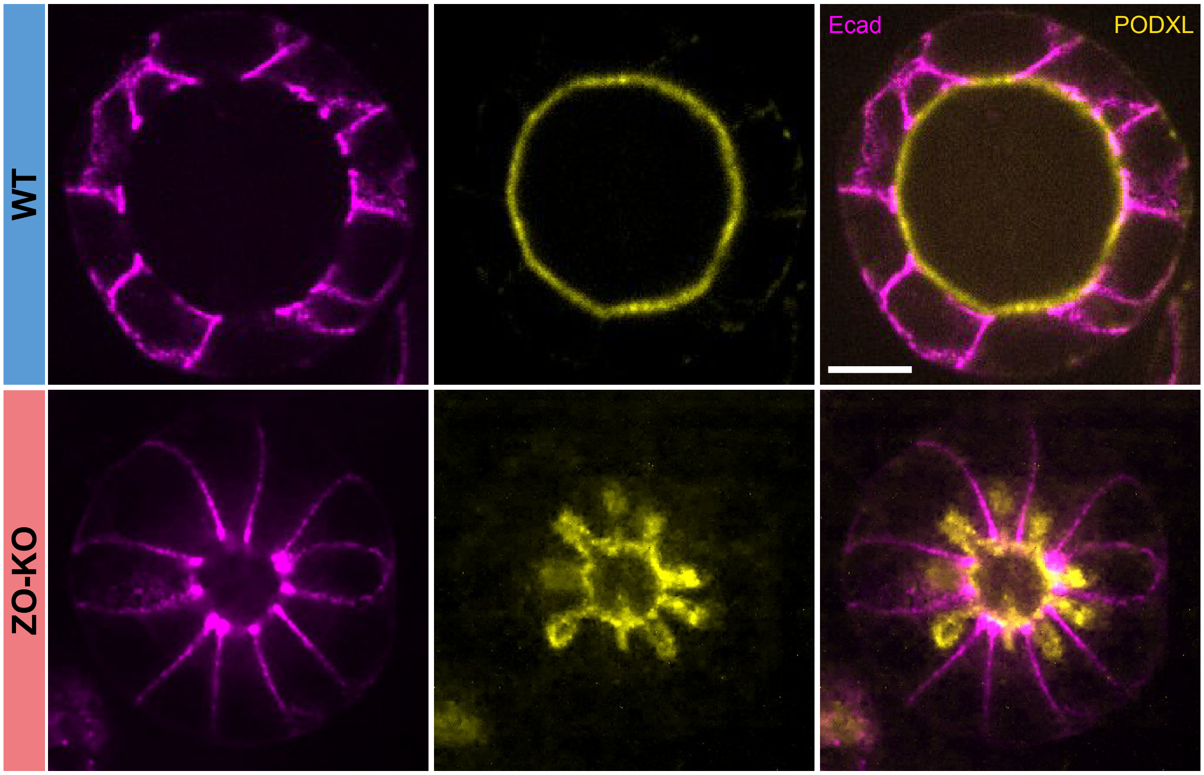


Supp. Info. Fig. 1: *Comparison between the MDCK WT vs MDCK ZO-KO lumen morphology - Spinning disk microscopy slice of MDCK WT spheroid - Spinning disk microscopy slice of MDCK ZO-KO spheroid - Lateral membranes are stained by E-cadherin (magenta) and apical membranes by Podocalyxin (yellow) - Scale bar: 10 μm*

and, by construction, the apical area of each cell is given by $S = S_{\text{paraboloid}}/N$. Then, the volume of the lumen is : $V = \frac{4}{3}\pi r_{\text{int}}^3 + \frac{N}{2}\pi r_a^2 h$ where $r_{\text{int}}$ is the radius of the sphere connecting all apical vertices.

The key assumption of our model is that the lumen morphology which minimizes the free energy of the lumen surface, $F$, is realized. The free energy is assumed to be given by the sum of three contributions; hydrostatic pressure, line tension along all the cell-cell junctions (apical junctional tension) and apical surface bending:

$$F = F_{\text{hydr. press.}} + F_{\text{apical junctional tension}} + F_{\text{apical surface bending}} \quad . \tag{1}$$

with

$$F_{\text{hydr. press.}} = -PV \tag{2}$$

$$F_{\text{apical junctional tension}} = TL \tag{3}$$

and

$$F_{\text{apical surface bending}} = N\kappa \iint \mathrm{d}A \cdot H(r)^2 \tag{4}$$

$H(r)$ is the local mean curvature elaborated in paragraph 2. For Eq. (4), we have assumed zero spontaneous curvature and considered only the lowest-order terms of curvatures *i.e* $O(H^2)$ and

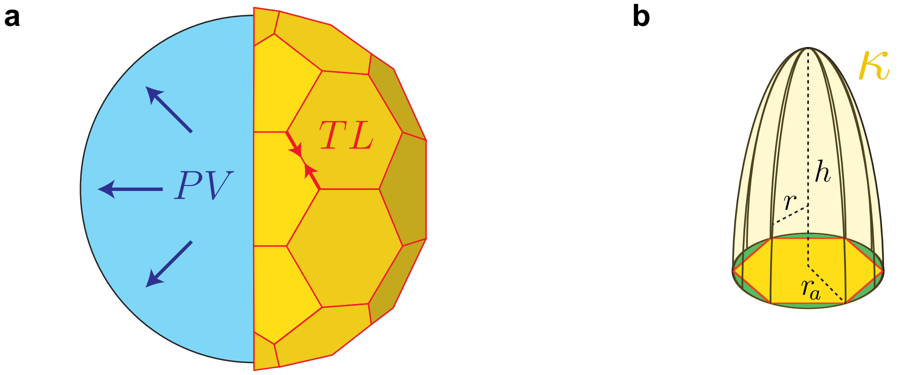


$$F = F_{\text{hydr. press.}} + F_{\text{apical junctional tension}} + F_{\text{apical surface bending}}$$

$$F = \quad -PV \quad + \quad TL \quad + \quad N\kappa \iint \mathrm{d}A \cdot H(r)^2$$

Supp. Info. Fig. 2: *Schematics of the simplified model of a spheroid and apical surface area and the different free energy contributions from hydrostatic pressure of the lumen, apical junctional line tension and apical surface bending - (**a**) The lumen surface is modeled as a tesselation of hexagons of total junctional length $L$ under the tension $T$ and surrounding a pressurised luminal volume $V$ - (**b**) Each cell apical surface is modeled by a paraboloid of based radius $r_a$ and height $h$ and bending rigidity $\kappa$.*

$O(K^1)$ with the Gaussian curvature $K$; the Gaussian-curvature term is omitted since $\int dAK$ is constant unless topology of the lumen surface changes due to Gauss-Bonnet Theorem [2]. The constant parameters $P$, $T$ and $\kappa$, refer to the difference of lumen hydrostatic pressure, the apical junctional tension and the bending modulus of the apical surface, respectively. We further assume that the minimization of F is given under the constraint that the total lumen surface $S$ should be splitted equally on each individual cell:

$$\frac{S}{N} = S_{\text{paraboloid}} = \frac{\pi r_a}{6h^2}\left[(r_a^2 + 4h^2)^{\frac{3}{2}} - r_a^3\right] \tag{5}$$

#### 2 First case: apical surfaces with no bending energy

We first consider the simpler case where the lumen morphology is determined by minimizing the free energy of the lumen surface, $F_1$.

$$F_1 = F_{\text{hydr. press.}} + F_{\text{apical junctional tension}} = -PV + TL \tag{6}$$

In this case, we can derive the existence of a transition line between two states: an inflated lumen or a fully collapsed lumen).

With the above parametrisation it is possible to compare the free energy of an observed flower lumen shape with a spherical lumen. If we set the free energy of the flower shape to 0:

$$F_{\text{spherical lumen}} - F_{\text{flower lumen}} = -P\frac{S^{\frac{3}{2}}}{6\sqrt{\pi}} + T\sqrt{2SN\sqrt{3}} \tag{7}$$

We have $F_{\text{spherical lumen}} < F_{\text{flower lumen}}$ (spherical lumen shape stable) if:

$$T < P\frac{S}{\sqrt{72\pi N\sqrt{3}}} \tag{8}$$

The line $T = P\frac{S}{\sqrt{72\pi N\sqrt{3}}}$ defines two regions in the $(P, T)$ phase diagram for lumen surface (see Supp. Info. Fig. 3a). If $T < P\frac{S}{\sqrt{72\pi N\sqrt{3}}}$, then $F_{\text{spherical lumen}} < F_{\text{flower lumen}}$ and spherical lumen is stable. Otherwise, the flower shape is stable.

### 3 Second case: apical surfaces with a bending energy

To compute the bending energy of the apical surface, one needs to express its mean curvature, with $r$, parametrizing the radius at height $z$, the paraboloid is parametrized as $z = S(r) = \frac{r^2}{2p}$, with $p = \frac{r_a^2}{2h}$. Then the mean curvature of this paraboloid can be computed as:

$$2H(r) = \frac{\frac{\partial^2 S}{\partial r^2}}{\left(1 + \left(\frac{\partial S}{\partial r}\right)^2\right)^{\frac{3}{2}}} + \frac{\frac{\partial S}{\partial r}}{r\left(1 + \left(\frac{\partial S}{\partial r}\right)^2\right)^{\frac{1}{2}}} \ , \tag{9}$$

so that

$$H(r) = \frac{1}{2p}\frac{\left(2 + \left(\frac{r}{p}\right)^2\right)}{\left(1 + \left(\frac{r}{p}\right)^2\right)^{\frac{3}{2}}} \tag{10}$$

We can now compute the free energy corresponding to the bending of the apical surface of cells:

$$\iint \mathrm{d}A \cdot H(r)^2 = \int_0^{2\pi} \mathrm{d}\theta \int_0^{r_a} \mathrm{d}r \left[ r\left(1 + \left(\frac{r}{p}\right)^2\right)^{\frac{1}{2}} \frac{1}{4p^2} \frac{\left(2 + \left(\frac{r}{p}\right)^2\right)^2}{\left(1 + \left(\frac{r}{p}\right)^2\right)^3} \right] \tag{11}$$

$$= \frac{\pi}{2}\left[\frac{4}{3} + \frac{1}{3p}\frac{3r_a^4 - 4p^4}{(r_a^2 + p^2)^{\frac{3}{2}}}\right] \tag{12}$$

$$= \frac{\pi}{6}\left[4 + \frac{\frac{6h}{r_a} - \frac{r_a^3}{2h^3}}{\left(1 + \left(\frac{r_a}{2h}\right)^2\right)^{\frac{3}{2}}}\right]. \tag{13}$$

Altogether, we obtain for the total free energy $F$:

$$F(r_a, h, P, T, \kappa, N) = -P\left[\frac{4}{3}\pi\left(\frac{N}{4}\right)^{\frac{3}{2}} r_a^3 + \frac{N}{2}\pi r_a^2 h\right] + TNr_a\sqrt{\pi\sqrt{6}} \tag{14}$$

$$+\frac{\pi N\kappa}{6}\left[4 + \frac{\frac{6h}{r_a} - \frac{r_a^3}{2h^3}}{\left(1 + \left(\frac{r_a}{2h}\right)^2\right)^{\frac{3}{2}}}\right] \tag{15}$$

In addition, for a given $r_a$, it is possible to compute the corresponding $h$ from Eq. (5), which can be rewritten as:

$$64h^4 + h^2 \left(48r_a^2 - \frac{36S^2}{\pi^2 N^2 r_a^2}\right) + 12\left(r_a^4 - \frac{Sr_a^2}{\pi N}\right) = 0 \tag{16}$$

In summary, one can compute $F(r_a, h, P, T, \kappa, N)$ from Eq. (15) with $h$ given as a solution of Eq. (16) and compute a phase diagram in the $(P, T)$ parameter space for the inner lumen radius (see Supp. Info. Fig. 3b).

Ranges of parameters are taken from experimental values (see Material and Methods) and are consistently used in Supp. Fig. 2 and in Fig. 5 with the vertex model.

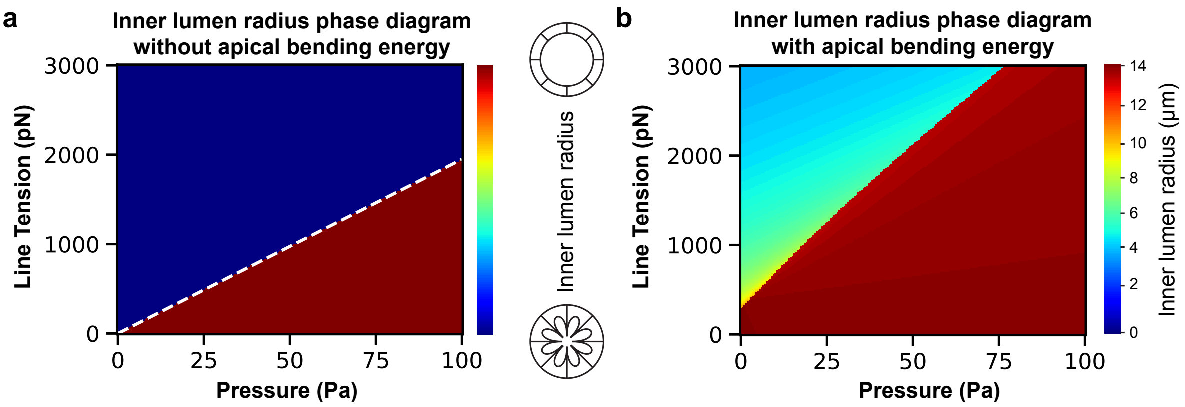


Supp. Info. Fig. 3: *Comparison of the phase diagrams of the inner radius of lumen determined by minimizing the free energy of the lumen surface without (**a**) and with (**b**) bending energy of the apical surface. Blue zones correspond to flower-like lumen shape and red zones to spherical inflated lumens. Total number of cells - $N = 42$ cells, total surface of lumen - $S = 2500$ $\mu m^2$, bending energy $\kappa = 2.10^{-16}$ J.*

### 4 Differences between the two phase diagrams

This linear relation described in paragraph 2 can be seen in Supp. Info. Fig. 3b with the addition of bending energy of the apical side, the slope of the transition line shifts towards higher values (Supp. Info. Fig. 3b). This analytical model also allows to probe the influence of the total number of cells and of the total lumen surface as illustrated in Supp. Info. Fig. 4. When the total apical surface $S$ increases the slope of the transition line scales up as well. Similarly, when the total number of cells $N$ in the cyst increases the slope of the transition line decreases. This is consistent with the estimation of the transition line from the simplified model obtained in section 2. The region of the phase diagram where the folded shape is more stable increases in size when $N$ increases. When $N$ increases, the inner lumen radius increases for a given set of $(P, T)$ values.

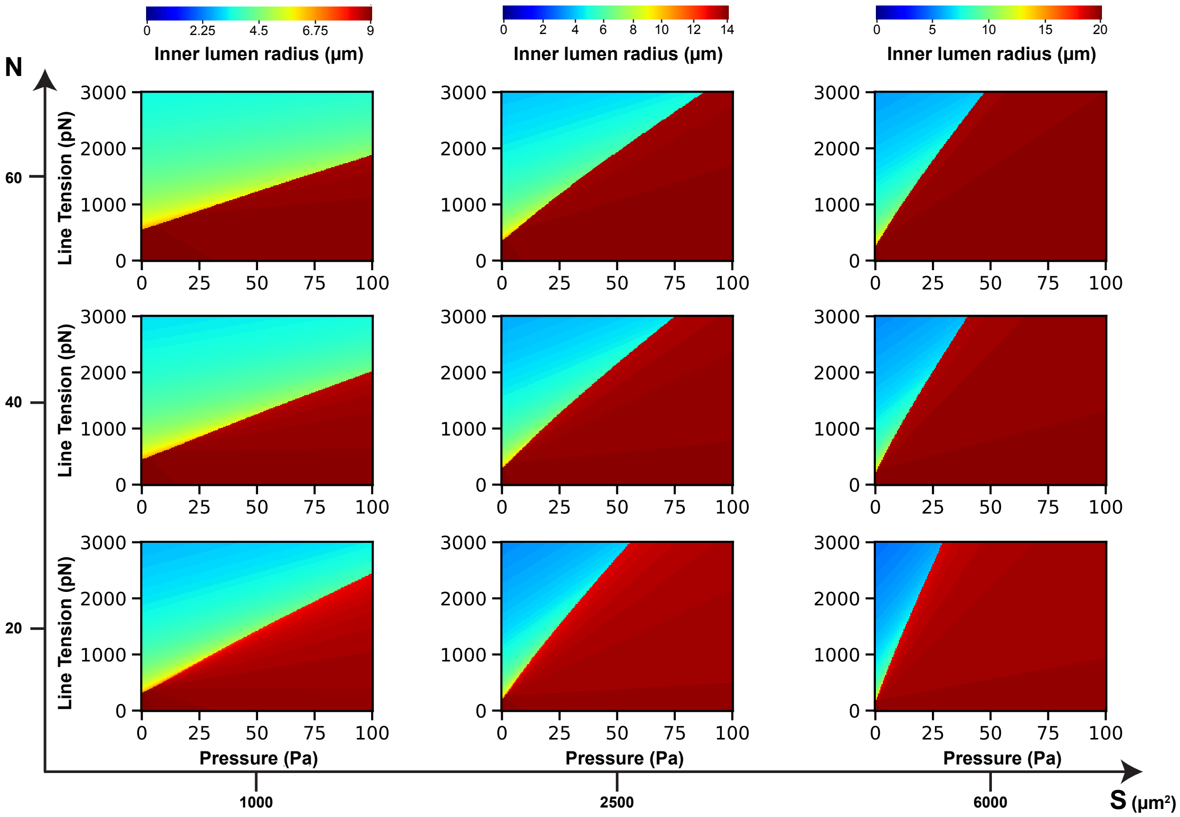


Supp. Info. Fig. 4: *Phase diagrams generated for (number of cells $N$ – total surface of lumen $S$) pair values for a bending energy -* $\kappa = 2.10^{-16}$ *J. Note that the inflated region (red zone) increases with larger $S$ values and decreases with larger $N$.*
